# Three-dimensional spatial quantitative analysis of cardiac lymphatics in the mouse heart

**DOI:** 10.1101/2023.02.01.526338

**Authors:** Evan H. Phillips, Vytautas P. Bindokas, Dahee Jung, Jay Teamer, Jan K. Kitajewski, R. John Solaro, Beata M. Wolska, Steve Seung-Young Lee

## Abstract

**Objective:** 3D microscopy and image data analysis are necessary for studying the morphology of cardiac lymphatic vessels (LyVs) and association with other cell types. We aimed to develop a methodology for 3D multiplexed lightsheet microscopy and highly sensitive and quantitative image analysis to identify pathological remodeling in the 3D morphology of LyVs in young adult mouse hearts with familial hypertrophic cardiomyopathy (HCM).

**Methods:** We developed a 3D lightsheet microscopy workflow providing a quick turn-around (as few as 5-6 days), multiplex fluorescence detection, and preservation of LyV structure and epitope markers. Hearts from non-transgenic (NTG) and transgenic (TG) HCM mice were arrested in diastole, retrograde perfused, immunolabeled, optically cleared, and imaged. We built an image processing pipeline to quantify LyV morphological parameters at the chamber and branch levels.

**Results:** Chamber-specific pathological alterations of LyVs were identified, but most significantly in the right atrium (RA). TG hearts had a higher volume fraction of ER-TR7^+^ fibroblasts and reticular fibers. In the RA, we found associations between ER-TR7^+^ volume fraction and both LyV segment density and median diameter.

**Conclusions:** This workflow and study enabled multi-scale analysis of pathological changes in cardiac LyVs of young adult mice, inviting ideas for research on LyVs in cardiac disease.

## 1. INTRODUCTION

The cardiac microenvironment includes nerves, immune cells, fibroblasts, extracellular matrix (ECM), blood vessels, and lymphatic vessels (LyVs). Among these components, the lymphatic system has been understudied despite its essential role in maintaining tissue fluid balance, interstitial pressure, and immune surveillance with drainage of inflammatory cytokines and antigens [1–3]. Compared to coronary arteries, less is known about the responses of cardiac LyVs in heart disease and heart failure [4]. Altered lymphatic function is a point of vulnerability in multiple cardiovascular disorders including hypertension, myocardial infarction, heart failure, and non-ischemic cardiomyopathies. With this evidence of lymphatic dysfunction and remodeling across a spectrum of cardiovascular disorders, there is a need for understanding how these impairments in the lymphatic system are linked to pathological processes [5]. To advance this understanding, we have developed an imaging methodology described here to determine morphological changes of cardiac LyVs in a pathological condition. To test our approaches, we have focused on the identification of structural changes of LyVs in the 3D cardiac microenvironment in a mouse model of familial hypertrophic cardiomyopathy (HCM). Late-stage HCM is characterized by significant interstitial fibrosis and lower microvessel density in the cardiac microenvironment [6,7]. Based on previous observations of remodeling in the cardiac microenvironment in a HCM genetic mouse model (mutated tropomyosin, Tm-E180G) [8,9], we could anticipate morphological alterations of LyVs in these hearts as well.

Immunofluorescence combined with fluorescence microscopy offers a high-resolution imaging assay for studying cardiac LyVs in mouse models of heart disease. In addition, recent reports have demonstrated the great potential of optical tissue clearing and lightsheet fluorescence microscopy (LSFM) for investigating cardiac LyVs [10,11]. However, at the same time, these previous works have also revealed technical challenges for 3D microscopy of cardiac LyVs, including achievement of sufficient heart transparency, multiplex immunolabeling for LyVs and other markers, and preservation of superficial LyV integrity. Further, a quantitative approach for comprehensive regional analysis of cardiac LyVs on the surface of the mouse heart has not yet been developed. To detect small and detailed morphological changes (or remodeling) of cardiac LyVs in the mouse heart, whether induced by an acute injury or a slowly developing chronic disease, it is necessary to develop a highly sensitive and quantitative image analysis protocol.

To address these challenges, we have developed a method incorporating the following: LSFM of intact superficial cardiac LyVs in optically cleared mouse hearts alongside another biomarker (ER-TR7^+^ fibroblasts and reticular fibers) and quantitative 3D analysis of the morphology of LyVs across ventricles and atria. After multiplex immunofluorescence, the whole mouse heart was made optically transparent by modifying an organic solvent (ethyl cinnamate, EtCi)-based tissue clearing method [12]. Through a custom image processing workflow in 3D image analysis software, we extracted quantitative multi-level data of the morphology and numbers of LyVs as well as the chamber-specific density of ER-TR7^+^ fibroblasts and reticular fibers.

Our imaging and analysis protocol allows one to produce comprehensive 3D maps of LyVs in mouse hearts as well as quantitative profiles of regional morphological changes of LyVs. This method could be applicable for a variety of experimental purposes, not the least of which are investigations of regional LyVs over the course of cardiac disease development and testing LyV-targeted therapy candidates in the diseased mouse heart.

## 2. MATERIALS AND METHODS

### 2.1. Ethical compliance and animal model, care, and maintenance

Experiments were approved by the Animal Care and Use Committee of the University of Illinois Chicago and in compliance with the Guide for the Care and Use of Laboratory Animals as adopted by the U.S. National Institutes of Health. Experiments employed male and female 40 day-old (P40) NTG and TG mice expressing a mutated HCM-linked tropomyosin (Tm-E180G) in the FVBN background strain, as previously described [13].

### 2.2. Materials and equipment checklist

A list of the chemicals and reagents, supplies, and equipment used in this study is presented in Table S1. This list also serves as a reference for those applying the protocol outlined below. Details of the IgG antibodies used in this study for immunolabeling are also included.

### 2.3. Protocol

All mouse hearts were prepared as described below in detail (steps 1-8).

#### 2.3.1. Mouse heart preparation (from perfusion to tissue clearing)

1. Individual mice were weighed and prepared for *in situ* retrograde cardiac perfusion by first heparinizing (1000 U/kg body weight i.p.) prior to anesthesia. Mice were deeply anesthetized with isoflurane (as determined by lack of toe pinch reflex). A partial thoracotomy was performed followed immediately by injection of KCl (5 mL) through the inferior vena cava to arrest the heart in diastole. Once the heart had stopped beating, the chest cavity was opened and the lungs moved to the animal’s right side to fully visualize the descending aorta (Figure S1A).
2. A shallow level of PBS was added to the dissecting tray to prevent introduction of air bubbles during subsequent perfusion of PBS, 2% PFA, and 1:1 red blood cell lysis buffer (RLB):RPMI cell culture media through the aorta. PBS (pH 7.4) was slowly injected superior to the diaphragm until fluids started to run clear. 2% PFA (5 mL) followed over 2 min at an injection site slightly proximal. 1:1 RLB:RPMI (5 mL over 2 min) was then injected at a third injection site further proximal. The heart (with the aortic root) was dissected, gently dabbed to remove excess moisture, and then weighed.
3. Hearts were sequentially processed (steps 3 to 7) at 4°C while shaking and protected from light. Importantly, hearts were left unembedded and only handled by the aortic root. First, samples were washed in PBS (pH 7.4, 20 min × 3) prior to light immersion fixation with 2% PFA (8x sample volume, 30 min) and repeat PBS washes (pH 7.4, 20 min × 3). If residual blood was still apparent on a heart, it was incubated in 1:1 RLB:RPMI (30 min) (Figure S1B) followed by a repeat PBS wash (pH 7.4, 20 min).
4. Each heart was permeabilized and blocked by immersion in RPMI + 0.3%v/v Triton X-100 (10 mL, 18 h).
5. Immunolabeling was carried out in two steps. A staining buffer (SB) of RPMI + 1%w/v BSA was used in each step followed by washes in PBS (pH 7.4, 30 min × 4). First, each heart was incubated in 1 mL of SB with a primary goat anti-LYVE1 antibody (1:50) for 24 h followed by washing.
6. Each heart was then incubated in 1 mL of SB with: 1) AF647-conjugated anti-goat IgG (1:250) as a fluorescent secondary antibody binding to the primary LYVE1 antibody; and, 2) DyL550-conjugated rat anti-ER-TR7 (1:25) as a fluorescent primary antibody. Antibody conjugation to DyL550 and purification were performed ahead of time (see Supporting Information). After 24 h, hearts were washed in PBS.
7. Hearts were dehydrated in an ascending gradient of pH-adjusted EtOH (pH 9) in deionized water (i.e., 20, 50, 70, and 99%). Each heart was placed in a full 5 mL microtube for each step (at least 30 min × 1, except 99% 30 min × 2).
8. Finally, each heart was moved to room temperature (at least 5 min) and transferred into a glass vial with pH-adjusted (pH 9) EtCi [12] (20 mL) + α-thioglycerol (60 μL). Hearts were kept at RT and protected from light, becoming optically transparent (due to refractive index (RI) matching) after a minimum of 30 minutes (Figure S1B and Figure S2).

#### 2.3.2. Lightsheet fluorescence microscopy

9. Hearts were imaged on an UltraMicroscope II with Zoom body configuration (LaVision) using a 2X objective (NA 0.5; 6 mm WD) with a RI-matching collar and dipping cap. Each unembedded sample was handled by the aortic root and mounted horizontally in a 1 cm-long tissue holder (LaVision) against a plastic screw at one end and a plate on the other. The entire heart volume was imaged in an EtCi-filled chamber using the following parameters: 561 and 637 nm excitation, 620/60 nm and 680/30 nm emission filters, left and right lightsheets with horizontal focusing (merge type: Contrast), 1x zoom, sheet NA 12 μm, sheet width 30%, exposure 25 ms, xy pixel size 3.02 μm, and z-step 5 μm. Multi-plane XY image data were 16-bit and in TIFF-OME format.

#### 2.3.3. Image processing and analysis

10. TIFF-OME files were loaded into Amira 3D Pro v2021.1 (FEI Company, now ThermoScientific) for 3D visualization, image processing, segmentation, and quantitative regional analysis. See Supporting Information for details on the following description of the image processing pipeline. Cardiac chamber volumes (excluding lumens and valves of left ventricle (LV) and right ventricle (RV) were measured after using the 3D lasso tool (with Intersect option) for efficient segmentation followed by refinement with the 2D lasso. LYVE1 channel data were pre-processed by 3D unsharp masking to enhance LyV edges prior to thresholding at a set level across samples. Isolated pixels were filtered out by running Remove Islands, which refined the final binarized data. ER-TR7 channel data were pre-processed using an Amira module based on the Frangi filter [14] to enhance ER-TR7^+^ fibroblasts and reticular fibers from nearby autofluorescence. Enhanced signals were thresholded at a set level and compared against raw data to verify accurate and full representation. An Amira module (Centerline Tree) based on TEASAR (Tree-structure Extraction Algorithm for Accurate and Robust Skeletons) [15] was used to extract the centerlines of LYVE1^+^ tubular signals. Vessel centerlines (‘skeletons’) were compared against LYVE1 raw data to verify accurate and full representation. LyV segment parameters and the signal volume of ER-TR7^+^ fibroblasts and reticular fibers were computed in each cardiac chamber for quantitative regional analysis. LyV segments not meeting the following three criteria were filtered out of all analyses and calculations: segment diameter (> 6.04 μm, i.e., greater than two times the lateral resolution to satisfy the Nyquist criterion [16]), segment length greater than 5 μm, and segment length greater than the diameter of the same segment. In figures showing LyV centerlines as 3D spatial graphs (Figure 2A, Figure 5A, Figure S4E, Figure S5), all unfiltered LyV segments are displayed. Volume fraction of ER-TR7^+^ fibroblasts and reticular fibers was computed by normalizing signal volume to chamber volume.

### 2.4. Statistics

Statistical analysis was carried out in OriginPro 2022. Chamber-specific individual measurements or group averages ± SEM are presented throughout the figures. Two-sample t-tests were used to compare the following differences between NTG and TG groups: chamber volumes and chamber average measurements of LyVs (segment density, average segment diameter, and average segment length). Differences between histograms for NTG and TG groups (showing branch-level distributions for LyV morphological parameters after binning) were tested by Mann Whitney U-tests (non-parametric) and exact *p*-values were calculated. The interquartile range (IQR) of these distributions was assessed between NTG and TG groups to determine the directionality of change for a given parameter. Two-way analysis of variance (ANOVA) was used to compare differences in ER-TR7^+^ volume fraction according to two factors (chamber and genotype). ER-TR7^+^ volume fraction values were plotted against corresponding chamber values for LyV segment density as well as median LyV segment diameter. Linear regression analyses were run between these two sets of parameters. Pearson’s correlation coefficients and *p*-values were computed for the regression fitting of values from specific chambers. For all tests, a *p*-value of less than 0.05 was considered statistically significant.

## 3. RESULTS

The objectives of this study were to 1) establish a method for 3D multiplexed fluorescence microscopy and quantitative morphological analysis of cardiac LyVs and 2) test whether this method discriminated LyV morphology and number in different regions of NTG and TG hearts.

### 3.1. Brief characterization of mice used

It has been previously shown that by P15 cardiac LyVs are fully developed in control hearts [17]. We have previously reported that TG HCM hearts start to develop morphological changes (i.e. increase in atrial size) and diastolic dysfunction as early as at P14 and by 8 weeks these mice show a severe hypertrophic phenotype [8]. In the experiments reported here, we used P40 NTG and TG HCM mice. Basic characterization of these mice is presented in Table S2.

### 3.2. Workflow for tissue processing, imaging, and image data analysis

We developed a workflow for 3D multiplex LSFM imaging of cardiac LyVs with optimized protocols of collection, processing, immunolabeling, and optical clearing of whole mouse hearts (Figure 1A). To harvest intact mouse hearts without blood clots, we first treated mice with heparin and performed *in situ* retrograde perfusion (Figure S1A) in the step 1 and 2. We further incubated the extracted hearts in RLB to completely remove heme in step 3, which helped decolorize and enhance transparency of the hearts after later EtCi-based clearing (Figure S1B). After multiplex immunofluorescence, we used graded ethanol dehydration followed by RI matching with EtCi [12] to render NTG and TG hearts optically transparent despite persistent pigmentation (Figure 1B, Figure S2A). Using a mouse heart mounted in a spectrophotometer, we measured an approximately 6-fold higher percentage of transmitted visible light after 30 min incubation in EtCi as compared to when the heart had just been fixed at the end of step 3 (Figure S2B). To protect superficial cardiac LyVs against any inadvertent damage, we only handled samples by the aortic root and kept them immersed in buffer or solvent at all possible times. For LSFM imaging, individual unembedded hearts were oriented and horizontally mounted in a sample holder to preserve superficial cardiac LyVs (Figure 1C). The processed hearts were scanned at high resolution for multiplex fluorescence detection of cardiac LyVs, ER-TR7^+^ fibroblasts and reticular fibers, and endogenous autofluorescence by LSFM (Figure 1D). While ER-TR7^+^ fibroblasts and reticular fibers were found throughout the optical sections of the heart, most LyVs were detected in the epicardium (Figure S3).

**FIGURE 1.**
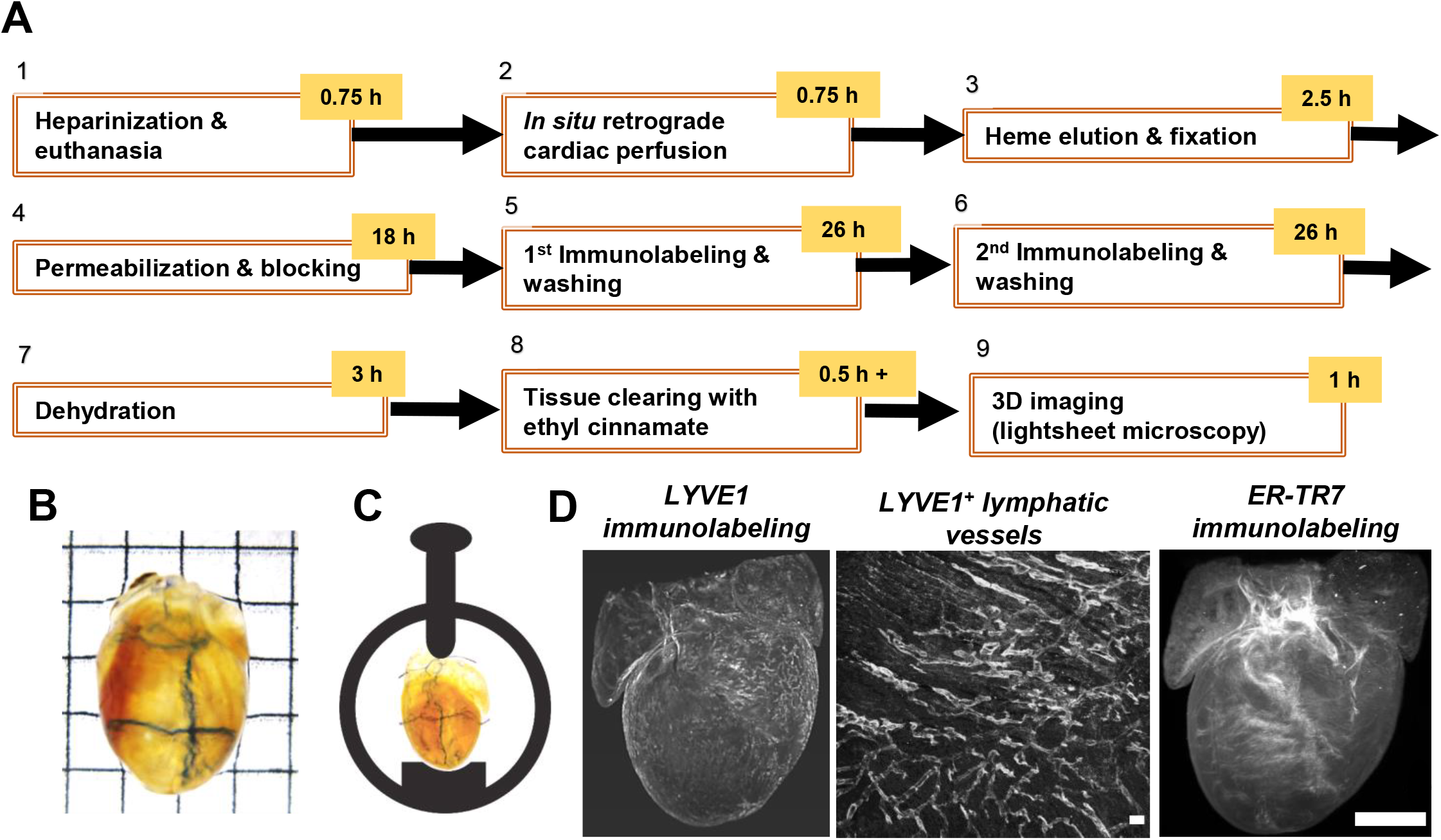
Workflow for multiplex lightsheet fluorescence microscopy. (A) Nine-step protocol to process and image whole mouse hearts. (B) Dehydrated and optically transparent whole P40 mouse heart used for imaging. Grid edge: 2.5 mm. (C) Orientation and horizontal mounting of heart in a sample holder. The heart is left unembedded and secured in place while immersed in ethyl cinnamate. (D) Left: Raw 3D image data of LYVE1 immunostaining. Middle: 2D maximum intensity projection showing LYVE1^+^ lymphatic vessels near ventricular surface. Scale bar: 0.1 mm. Right: Raw 3D image data of ER-TR7 immunostaining against endogenous autofluorescence. Scale bar: 1 mm.

**FIGURE 2.**
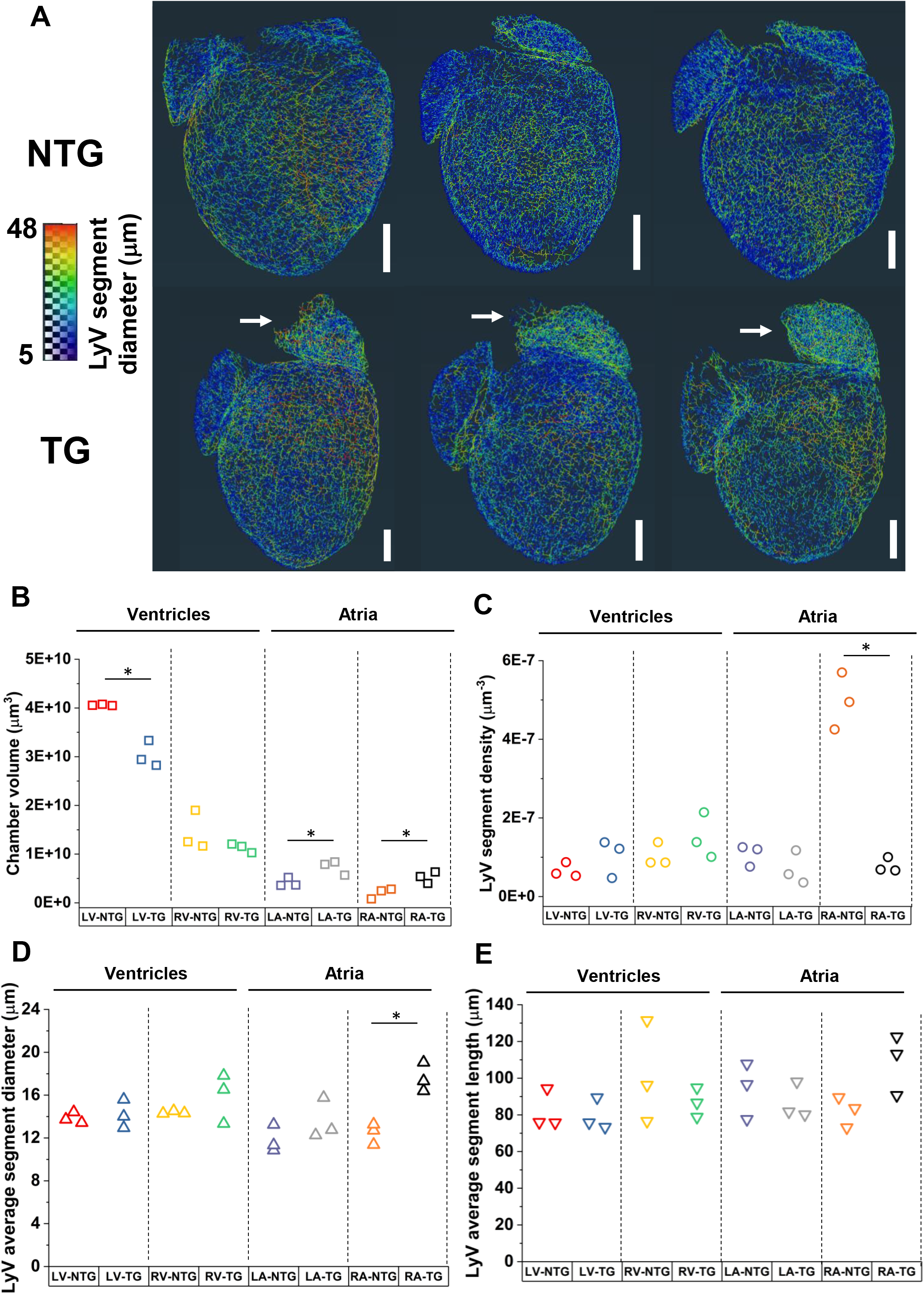
LyV distribution in NTG and TG P40 hearts and chamber-level measurements. (A) Superficial LyV segments are displayed as vessel centerlines and viewed in 3D from a posterior perspective of each imaged sample. The color scale corresponds to average diameter of a given LyV segment, with red being largest (48 μm maximum). Of note are the enlarged left and right atria in the TG hearts (white arrows point to RA). Scale bar: 1 mm. (B) Volume (μm^3^) of each chamber (LV and RV lumens excluded) in NTG and TG hearts was determined by 3D segmentation. (C-E) Chamber average values of the following parameters are plotted for each heart: density of LyV segments (C), LyV average segment diameter (D), and LyV average segment length (E). Note that number of LyV segments have been normalized to the respective chamber volume in (C). *, *p*<0.05 by two sample t-test; otherwise, no statistically significant difference between NTG/TG pairs for each chamber.

For quantitative 3D spatial analysis, we built an image processing workflow using Amira 3D Pro (Figure S4). We segmented the cardiac chamber walls, LyVs, and ER-TR7^+^ fibroblasts and reticular fibers using endogenous and exogenous fluorescence signals in 3D LSFM image data. The segmentation quality of ER-TR7^+^ fibroblasts and fibers and LYVE1^+^ LyVs were verified by comparison against raw data signal (Figure S4C and Figure S5 (left column), respectively). Centerlines of LYVE1^+^ tubes were extracted by skeletonization and the morphological parameters of the extracted LyVs (diameter, length, volume, and tortuosity) were projected in 3D and quantitatively analyzed after filtering (Figure S4D, E and Figure S5 (right column)). Furthermore, distribution and morphological patterns of LyVs and ER-TR7^+^ fibroblasts and reticular fibers and their spatial relationship were computed in each cardiac chamber for quantitative regional analysis (Video S1).

### 3.3. Regional differences in lymphatic vessel parameters

After running the custom-built image processing pipeline, we mapped the 3D distribution and size range of LyVs in NTG and TG hearts (Figure 2A). LyV segments are represented as vessel centerlines and are visible around the entire 3D surface of the heart with the color coding corresponding to the average diameter of a given LyV segment. We observed larger atria in TG hearts as well as wider LyVs (red colored) on the right side of TG hearts. We segmented the four cardiac chambers of each heart to compute chamber volumes (lumens excluded). We analyzed four parameters that summarize the collective size and number of LyV segments in the chambers of each heart: chamber volume, number of LyV segments per chamber volume, average segment diameter, and average segment length (Figure 2B-E). In TG hearts, LV volumes were significantly smaller while LA and RA volumes were significantly larger than those of NTG hearts (Figure 2B). Despite the difference in LV volumes, there was not a significant difference in LyV segment density between groups (Figure 2C). Interestingly, we found significantly fewer LyV segments per volume in the RA of TG hearts (Figure 2C) but also significantly larger average LyV segment diameter (Figure 2D). Other group comparisons did not show a significant difference in LyV segment density or average segment diameter. Average segment length was also not significantly different between NTG and TG hearts in any chamber (Figure 2E).

### 3.4. Branch-level measurements of lymphatic vessel parameters

We further analyzed the branch-level distributions of the following morphological parameters for LyV segments: diameter, length, volume, and tortuosity in the four chambers of NTG and TG hearts (Figure 3 and Figure S6). All distributions of LyV segment morphological parameters were non-normal and right-skewed. With respect to LyV diameter, the majority of LyV segments were smaller than 10 μm wide (Figure 3A, C, E, G). Smaller than 10 μm-wide LyVs segments are expected to be enriched in lymphatic capillaries while ones greater than 10 μm-wide are expected to be enriched in collector and pre-collector vessels with open lumens [18,19]. As compared to NTG hearts, fewer LyV segments less than 10 μm wide in all TG heart chambers were found, suggesting LyV rarefaction (Figure 3A, C, E, G). A greater number of wide (>14 μm) LyV segments presented in RA of TG hearts (Figure 3A). In RV and LV, fewer LyV segments in specific diameter ranges over 10 μm were found (Figure 3E, G). For LyV segment length, a greater number of long (>125 μm) LyV segments was seen in RA of TG hearts than NTG hearts (Figure 3B). In contrast, fewer LyVs segments in other chambers of TG hearts were in the same length range (Figure 3D, F, H). We summarize these observation in Table 1 after analyzing the interquartile ranges (IQRs) and running statistical tests. In brief, there was a statistically significant increase in the number of LyVs wider than 14 μm (2.2-fold greater) and longer than 125 μm (2.3-fold greater) in RA of TG hearts (Table 1). Further, we noted the following trends for LyVs in TG hearts: 1) For segment diameter, fewer LyVs less than 10 μm wide across all chambers and fewer 10-32 μm wide in LV; 2) For segment length, fewer longer than 100 μm, 150 μm, and 125 μm in LA, RV, and LV, respectively.

**FIGURE 3.**
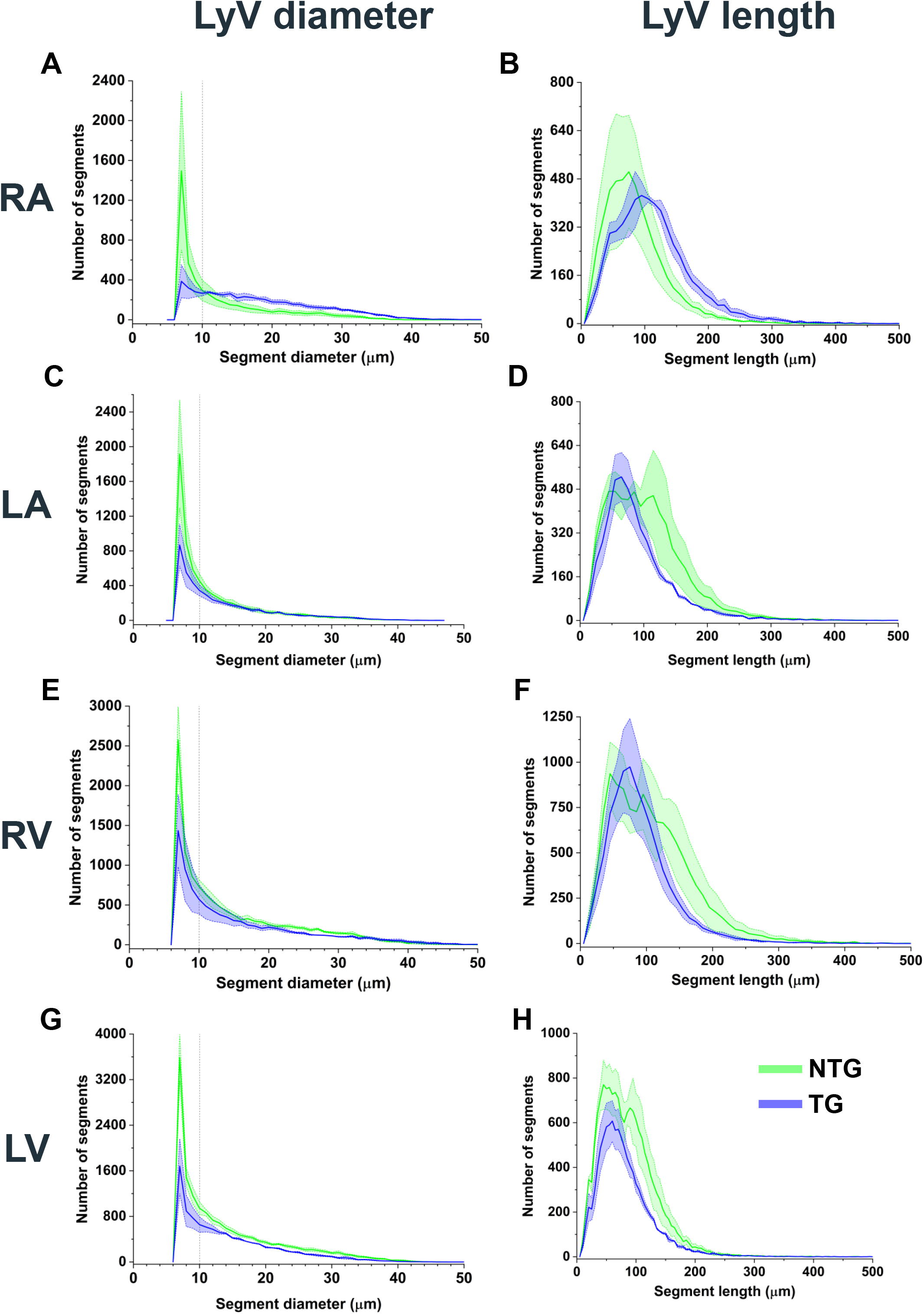
Branch-level measurements of LyV segments across chambers. Histograms show distributions of LyV segments for NTG (green) and TG (blue) hearts according to segment diameter (left column: A, C, E, G) and segment length (right column: B, D, F, H) in RA, LA, RV, and LV. Bin sizes: 1 μm (diameter) and 10 μm (length). Thick distribution lines: average. Shaded areas: standard error of mean. Vertical dotted lines (A, C, E, and G) mark 10 μm diameter cutoff. Interquartile ranges were analyzed and Mann Whitney U-tests were run to compare each set of binned distributions for NTG and TG hearts (see Table 1).

**FIGURE 4.**
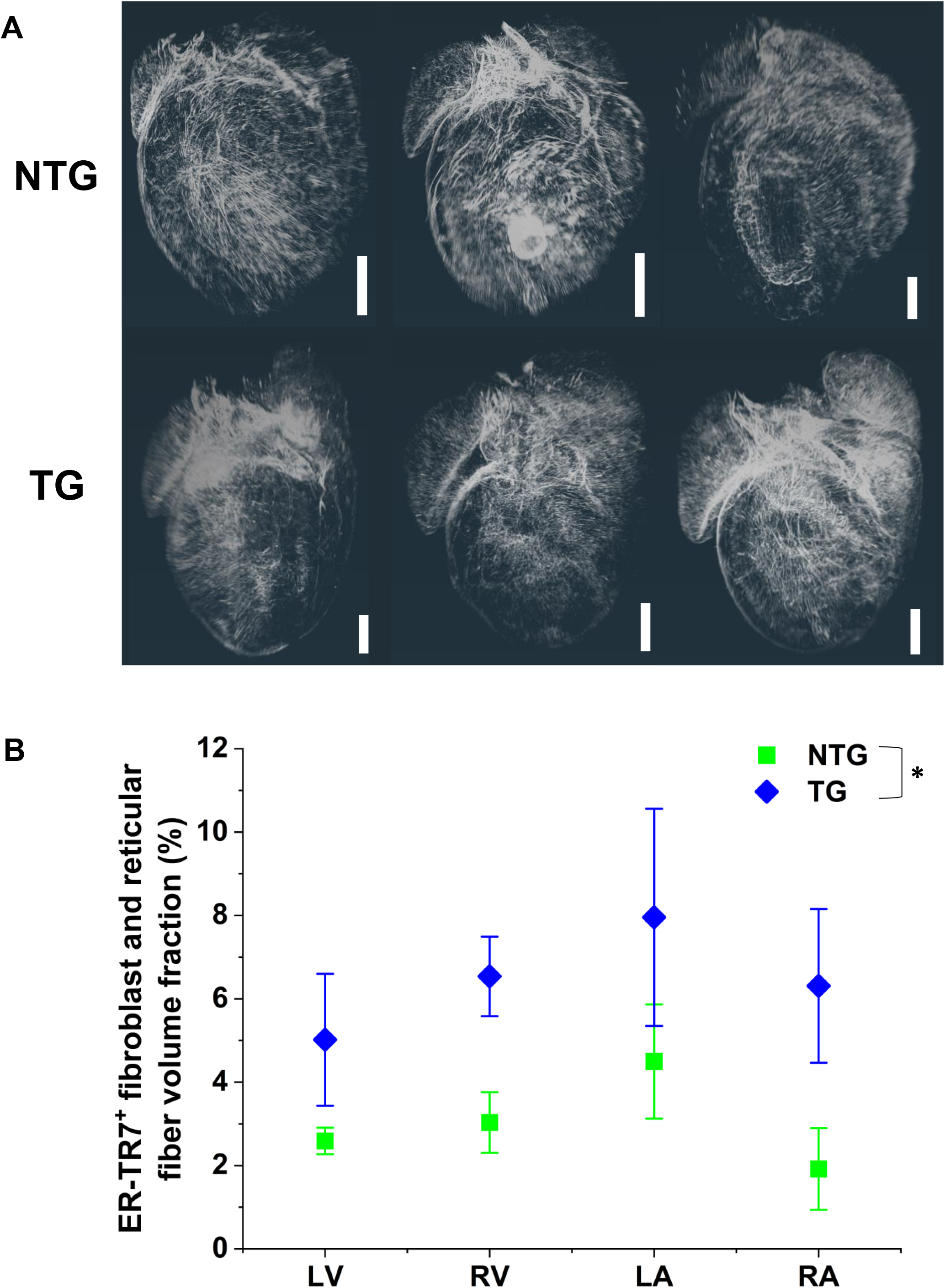
Distribution and chamber-level measurements of ER-TR7^+^ fibroblasts and reticular fibers. (A) Voxelized renderings of the final structure-enhanced ER-TR7^+^ fibroblasts and reticular fibers in all chambers are shown in 3D from a posterior perspective of each imaged sample. Scale bar: 1 mm. (B) Volumetric fraction of ER-TR7^+^ fibroblasts and reticular fibers measured in individual chambers of NTG and TG hearts. There was a statistically significant difference in the population means of the NTG and TG groups (two-way ANOVA; *, *p*=0.0042).

**FIGURE 5.**
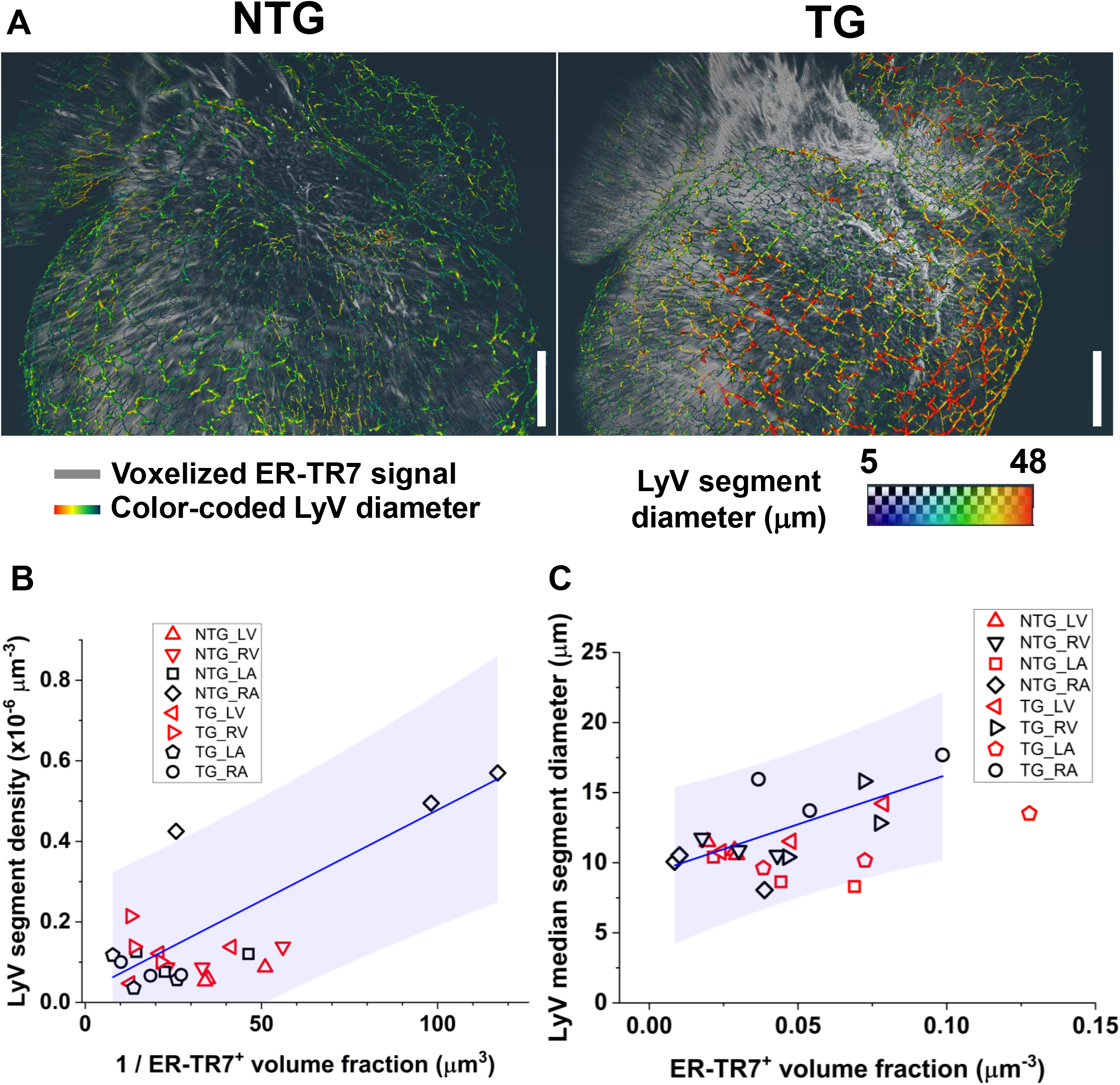
Association between ER-TR7^+^ volume fraction and LyVs. (A) LyVs tubes (color-coded according to segment diameter) overlaid on voxelized fibroblasts and reticular fibers (grayscale) from a posterior view of the top half of an NTG (left) and TG (right) heart. Scale bar: 700 μm. The color scale corresponds to average diameter of a given LyV segment, with red being largest (48 μm maximum). (B) Inverse association between ER-TR7^+^ volume fraction (Figure 4B) and LyV segment density (Figure 2C) in RA and LA (black colored symbols) in TG and NTG hearts. RV and LV values (red colored symbols) failed to fit linear regression curve. (C) Association between ER-TR7^+^ volume fraction (Figure 4B) and LyV median segment diameter in RV and RA (black colored symbols) in TG and NTG hearts. LV and LA values (red colored symbols) failed to fit linear regression curve. Blue shaded areas represent the 95% prediction interval for the regression curves.

**TABLE 1.**
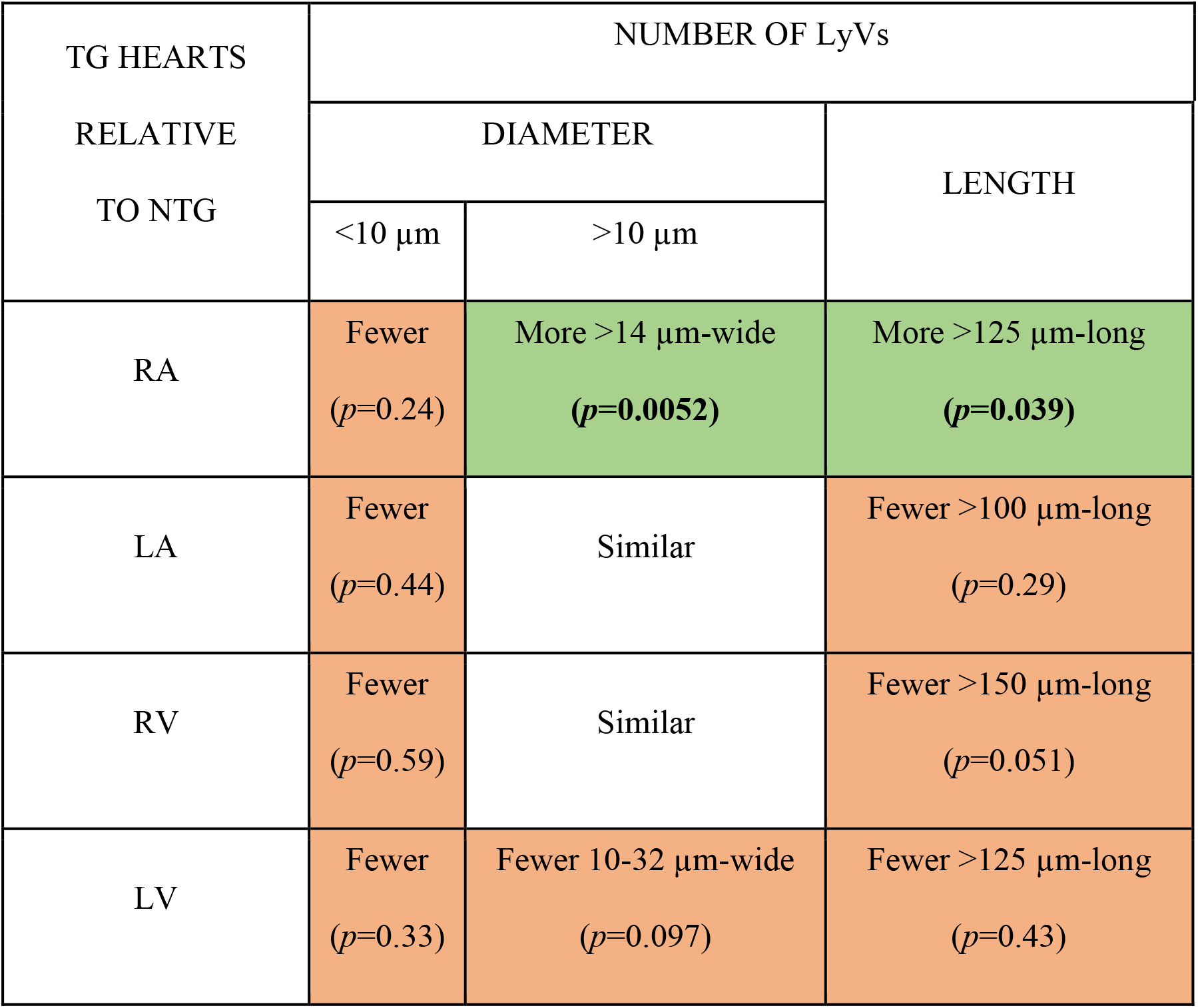
Branch-level summary of observations from binned distributions of LyV segment parameters. LyV segment diameters (less and greater than 10 μm) and lengths were compared across the interquartile ranges for TG hearts against the range of values for NTG hearts. Binned distributions were also statistically compared by Mann-Whitney U-tests (exact *p*-values are shown). Green-and orange-shaded cells indicate an increase and decrease, respectively, relative to NTG hearts. Unshaded cells indicate a lack of consistent increase or decrease over the range of values. Bolded *p*-values indicate statistical significance (*p*<0.05). See histograms in Figure 3 for binned whole distributions.

| TG HEARTS<br>RELATIVE<br>TO NTG | NUMBER OF LyVs |  |  |
| --- | --- | --- | --- |
|  | DIAMETER |  | LENGTH |
| | <10 $\mu\text{m}$ | >10 $\mu\text{m}$ | |
| RA | Fewer<br>($p=0.24$) | More >14 $\mu\text{m}$-wide<br>**($p=0.0052$)** | More >125 $\mu\text{m}$-long<br>**($p=0.039$)** |
| LA | Fewer<br>($p=0.44$) | Similar | Fewer >100 $\mu\text{m}$-long<br>($p=0.29$) |
| RV | Fewer<br>($p=0.59$) | Similar | Fewer >150 $\mu\text{m}$-long<br>($p=0.051$) |
| LV | Fewer<br>($p=0.33$) | Fewer 10-32 $\mu\text{m}$-wide<br>($p=0.097$) | Fewer >125 $\mu\text{m}$-long<br>($p=0.43$) |

Differences seen in LyV segment diameter scaled with LyV segment volume (Figure S6A, C, E, G). In RA of TG hearts compared to NTG hearts, there was a higher median tortuosity in LyV segments with less than 10 μm wide (1.26 vs 1.17; *p*=0.36) and ones more than 10 μm wide (1.19 vs 1.15; *p*=0.39) (Figure S6B). Differences in tortuosity varied depending on the LyV size in LA, RV, and LV (Figure S6D, F, H). After branch-level comparisons to NTG hearts, we identified distinct morphological changes of LyVs in RA of TG hearts.

### 3.5. Group differences in fibroblasts and reticular fibers and association with lymphatic vessel parameters

ER-TR7 was selected as a second marker to evaluate structural cell type-specific density changes in TG hearts. ER-TR7 is a cytoplasmic antigen found in fibroblasts but also in the ECM as a secreted component [20]. ER-TR7 immunolabeling has previously been studied in the heart [21]. Qualitatively, we did not see a difference in the density of ER-TR7^+^ signal between groups (Figure 4A). Measurements in individual chambers of TG hearts all showed an increase relative to NTG hearts (Figure 4B): LV (1.9 ± 0.65 fold); RV (2.2 ± 0.61 fold); LA (1.8 ± 0.79 fold); and, RA (3.3 ± 1.9 fold). None of these pairwise differences was significant, but there was a statistically significant difference between the NTG and TG population means (*p*=0.0042).

We further investigated whether there was an association between ER-TR7^+^ volume fraction (Figure 4) and LyV segment density (Figure 2) at the chamber level. Despite the range in LyV segment diameters, we also looked at whether larger median LyV diameter correlated with higher ER-TR7^+^ volume fraction. Figure 5A provides 3D image data from an NTG and TG heart illustrating these observations. RA and LA measurements of LyV segment density for all NTG and TG hearts were found to be inversely associated with ER-TR7^+^ volume fraction (Figure 5B; Pearson’s r=0.84; *p*=7.1 × 10^−4^). RV and RA measurements of LyV median segment diameter for all NTG and TG hearts were found to be associated with ER-TR7^+^ volume fraction (Figure 5C; Pearson’s r=0.67; *p*=0.017). These results suggest further investigation of the association between changing morphology and density of LyVs and ER-TR7^+^ structural cells and stromal fibers in this HCM model.

## 4. DISCUSSION

### 4.1. Motivation to develop 3D immunolabeling and optical tissue clearing protocol

Here we present a 3D immunolabeling and clearing protocol allowing for preservation and quantification of LyV morphology alongside at least one other marker. In addition, we developed an image processing pipeline for whole mouse hearts that enables regional quantitative analysis of LyV morphology and volumetric density of additional markers. The cardiac microenvironment is full of complex 3D structures including myofibers, connective tissue, and blood and lymphatic vessel networks. However, *ex vivo* analysis of the heart is generally carried out by 2D thin-section immunohistochemistry or immunofluorescence whereby cellular markers and morphology are visualized in cross-section. As a result, cross-sections (typically less than 10 μm thick) provide just a small snapshot of processes, such as vascular remodeling, without depth resolution. Furthermore, lymphatic capillaries are fragile thin-walled networks composed of endothelial cells with a range of diameters (due to their distensibility). For these reasons, 3D image analysis is necessary to provide not only actual measurements but also comprehensive and unbiased analysis of morphological features such as vessel density, branching, and associations among different cell types.

### 4.2. Workflow testing and advantages

In developing this workflow, we compared previously published 3D immunolabeling and clearing protocols to identify factors that are either detrimental or ones that can be omitted to shorten a protocol while still allowing for detection and regional quantification of multiplex fluorescence in the mature mouse heart. Our workflow requires as few as five to six days to prepare the tissue followed by imaging for an hour and semi-automated image processing for two or more hours. Some clearing techniques require the use of chemicals that may damage or crosslink the epitopes of important cellular markers [22,23]. We ran multiple preliminary tests with different chemicals, such as m-xylylenediamine (MXDA), which was reported by Zhu et al. [24] to decolorize heme-rich tissues and enhanced optical transparency in mouse embryos. We tested the protocol by incubating a mouse heart in increasing concentrations of sorbitol and MXDA. Although the method provided exceptionally good decolorization and optical transparency in young adult mouse hearts, LyVs could not be detected with immunofluorescence after MXDA incubation testing (data not shown), likely due to the loss of structural integrity and antigenicity of LYVE1 epitopes.

In developing an optimized protocol and workflow, we applied light fixation [25] to retain antigenicity of easily cross-linked epitopes and modified a cardiac-specific protocol [12] by Merz *et al*. that recommended hydrogen peroxide and EtCi for autofluorescence homogenization and optical tissue clearing, respectively. The trade-off we found for hydrogen peroxide is that while it may enable better resolution of cardiomyocytes [12] or the structure of single cells, there is a risk of artificially dilating lymphatic vessels (due to high peroxidase content) and quenching existing fluorescence (e.g. fluorescent protein). Nonetheless, the endogenous autofluorescence of the heart can be both exploited and overcome. By using bright and photostable fluorophores and making use of image processing (e.g. Frangi filter [14,26]), we could discriminate fibroblasts and reticular fibers as well as LyVs despite underlying autofluorescence in each wavelength channel. Thus, our results suggest that hydrogen peroxide can be omitted. For the purpose of cardiac chamber segmentation, the signal from endogenous autofluorescence without bleaching was sufficient to visualize differences in cardiac muscle and connective tissue structure. pH adjustment of ethanol and EtCi to pH 9 ensured that our samples retained fluorescence signals for at least many months after immunolabeling and clearing. As well, addition of the reducing agent α-thioglycerol prevented gradual loss of optical transparency.

To enhance optical transparency of mouse hearts, we applied RLB for heme elution. *In vivo* heparinization followed by retrograde perfusion with PBS alone and then sequential incubations in RPMI with Triton X-100 and BSA, an ascending ethanol gradient, and pH-adjusted EtCi caused a heart to mildly decolorize and achieve a moderate optical transparency after incubation in EtCi due to the unremoved hematogenous pigments. However, pre-processing with RLB helped to remove heme and then enhance optical transparency of mouse hearts after EtCi-based clearing process. Taken together, these tests provide evidence for the necessary reagents (RLB, RPMI, BSA, Triton X-100, EtOH, and EtCi) to enable preservation and detection of LyVs in young adult hearts while making the heart optically transparent for LSFM.

Multiplex fluorescence detection was also a key consideration in the development of this workflow. We used the longest excitation wavelengths available to us (561 and 637 nm) to detect autofluorescence and antibody markers in each channel. Due to more limited depth penetration, shorter wavelengths were not useful in our hands for whole heart imaging. The high autofluorescence of the heart (and other organs) seen in a shorter wavelength channel, such as 488 nm, may be detrimental or helpful. A near-infrared channel (e.g. 785 nm) on the other hand is available for some LSFM users and would be beneficial for further multiplexing.

We built a semi-automated image processing pipeline that handles large multi-channel 3D image data and outputs regional measurements of morphological parameters and volumetric density. We implemented an Amira module (Centerline Tree) that is well-suited to detecting the tree-like structure of LyVs (i.e. large collecting vessels down to blunt-ended capillaries without loops). We used the autofluorescence in the lower wavelength channel (561 nm), which provided sufficient detail through the depth of the hearts to manually but efficiently segment each chamber wall in 3D. Ultimately, we acquired regional distributions for multiple parameters. The distributions represented both closed-lumen and open-lumen LyVs. The latter included dilated LyV capillaries, pre-collectors, and collecting vessels, all of which are known to be present in the mouse heart subepicardium and outer myocardium [11,27].

### 4.3. Lymphatic vessel changes and HCM

We acquired novel data indicating that morphological changes in LyVs occur in a transgenic model of HCM alongside changes in the volumetric density of fibroblasts and reticular fibers. Interestingly, we observed the most distinct differences in LyV segment density, diameter, and length in the RA of TG hearts. Larger median segment diameter, longer median segment length, and higher median segment tortuosity were noted across LyV segments in the RA as compared to the RA in NTG controls. When normalized to the larger chamber volumes, we found that the density of LyV segments in RA was significantly decreased, opposite to the effect for ER-TR7^+^ fibroblasts and reticular fibers. We also found that higher median LyV diameter in RV and RA correlated with higher ER-TR7^+^ volume fraction. We have reported that the atria do expand in size very early in Tm-E180G TG development and large thrombi are known to form in the atria [13]. Remodeling and/or flow disruption of coronary microvessels may therefore be related to chamber-specific morphological changes in LyVs. Whatever the case, our results suggest the possibility of altered LyV function in interstitial fluid balance and clearance of immune cells in HCM with specificity to the RA. Although it has been known for many years that there is a role for the immune system in heart failure, including HCM [28], little is known regarding modifications in LyVs in these processes. Anatomical studies have reported a lower abundance of adult mouse cardiac LyVs in atria versus ventricles [27]. However, there remains a lack of information and understanding regarding atrial LyVs. Nevertheless, it is apparent that our finding of a significant modification in LyV morphology in the atrium may have functional consequences related to cardiac edema with elevated interstitial compression of myocytes and clearance of immune cells, cytokines, and antigens [4,19]. Moreover, modifications of the RA known to be causal in induction of atrial fibrillation would be expected to be exacerbated with modifications in lymphatic function [29]. In fact, there is evidence of right ventricular and right atrial fibrillation in a subset of HCM patients [30]. Given the paucity of literature on atrial LyVs, further investigation is necessary to interpret the significance of LyV morphological changes in HCM.

### 4.4. Future research

The development of an advanced imaging method for cardiac LyV research in the mature mouse heart opens the door to new research questions. The ultimate advantages that come out of applying this method include fluorescence multiplexing, chamber-specific regional quantification, and LyV segment distribution analysis. These aspects motivate analysis of LyVs in cardiac models alongside investigation of the cardiac immune microenvironment, fibrosis, and microvessel rarefaction. Pharmacologically stimulating LyV growth may also be beneficial in contexts outside of myocardial infarction [19] and edema [31], such as alleviating fibrosis [32,33]. Furthermore, omics analysis [34,35] provides complementary hypothesis-generating questions about the diverse cell populations in the heart, which may be best tested with 3D multiplex imaging.

## 5. PERSPECTIVES

- Our workflow with optimized protocols enables 3D multiplex lightsheet microscopy of cardiac lymphatic vessels in whole mouse hearts.
- Lower density and larger diameter of cardiac lymphatic vessels in RA of HCM TG mouse hearts were found compared to RA of NTG mouse hearts.
- Density and median diameter of lymphatics vessels in RA were also found to be associated with the volume fraction of fibroblasts and reticular fibers.

## LIST OF ABBREVIATIONS

ECM: Extracellular matrix
EtCi: Ethyl cinnamate
HCM: Hypertrophic cardiomyopathy
IQR: Interquartile range
LSFM: Lightsheet fluorescence microscopy
LyVs: Lymphatic vessels
NTG: Non-transgenic
RLB: Red blood cell lysis buffer
RI: Refractive index
3S: Three-dimensional
TG: Transgenic
LV and LA: Left ventricle and atrium
RV and RA: Right ventricle and atrium

## ACKNOWLEDGEMENTS

We thank Koreena Rafael-Clyke for mouse colony maintenance and genotyping (University of Illinois), Northwestern University Center for Advanced Microscopy for access to LaVision UltraMicroscope II, and Ebba Brakenhielm (INSERM) and Xiaolei Liu (Northwestern University) for helpful discussions.

## AUTHOR CONTRIBUTIONS

E.H.P., J.K.K., R.J.S., B.M.W., and S.S.-Y.L designed the study. E.H.P. performed collection, processing, and imaging of moue hearts. E.H.P. and V.P.B. developed image processing workflows and analyzed image data. D.J. and J.T. processed and took optical measurements of moue hearts. E.H.P., D.J., and J.T. conducted data analysis. E.H.P., R.J.S., B.M.W., and S.S.-Y.L. wrote and edited the manuscript. All authors approved of the final manuscript.

## SUPPORTING INFORMATION

Additional supporting information can be found in the online version of the article at the publisher’s website.

