## Supporting Information for "Three-dimensional spatial quantitative analysis of cardiac lymphatics in the mouse heart"

**Antibody conjugation and dialysis**

Monoclonal rat anti-mouse ER-TR7 antibody (BioXCell) was conjugated to an amine-reactive fluorescent dye (DyLight 550 NHS Ester, Thermo Fisher) and dialyzed in-lab. The following steps were carried out at 4°C for the times indicated below.

**Antibody-dye conjugation**

- Dilute stock antibody (2.92 mg/mL) to 1.5 mg/mL in 1x PBS (pH 8.0)
- Add 20 molar-fold excess DyL550 (10 mg/mL in DMF) to diluted antibody solution
- Incubate on a shaker with light protection (18 h)

**Purification**

- Hydrate a dialysis cassette (10,000 MWCO Slide-A-Lyzer cassette, Thermo Fisher) in PBS (10 min)
- Follow product instructions to inject the fluorescent dye-conjugated antibody into the dialysis cassette using an appropriately sized syringe with 18G needle
- Allow solution to dialyze (overnight) in a light-protected 2 L beaker filled with PBS while rotating on a magnetic stir plate
- Chang the dialysis buffer the following morning and once again in the early evening to ensure complete removal of unreacted dye from the dialysis cassette
- Follow product instructions to remove the fluorescent dye-conjugated antibody from the dialysis cassette using an appropriately sized syringe with 18G needle
- Store DyL550-conjugated ER-TR7 antibody at 4°C
- Perform immunolabeling with titrated antibody during subsequent 3 months

**Image processing pipeline**

A custom image processing pipeline in Amira 3D Pro (FEI company) is detailed below.

Cardiac chamber segmentation and mask generation, chamber wall quantification, semi-automated image processing of fluorescence signals, and quantitative analysis are the major steps.

The images and data presented in the manuscript were generated using this pipeline for every sample. Any underlined text indicates key values or options, which were chosen to generate the presented results. Other users are suggested to test these for their own data.

**Cardiac chamber segmentation and mask generation**

Load 16-bit TIFF-OME raw image data for the fluorescence channel that best shows autofluorescence of the heart muscle (e.g. 561 nm)

- Click the data object in the Project Viewer and switch to the Segmentation Editor (Classic workroom)
- Use the Thresholding tool (7^th^ tab in selection window) to select all pixels corresponding to the heart (i.e., all pixels that are not background)
- Use the masking slider bar to set a lower limit for intensity (e.g. 1200)
- Verify that the mask captures the edges of the heart across all slices
- The mask does not need to fill all inside holes
- Click ‘Select Masked Voxels’
- Click ‘Add’ under materials and create a new material called “All signal”
- Highlight “All signal” (left-click the name)
- Click the ⊕ sign (top of selection window) to assign the selected pixels to the material
- Click the ‘Select’ button to the right of the material and checkmark ‘3D’ for the material
- This will make the pixels in the material an active selection
- In the 3D viewer, you should see the heart surface

*As you go through the steps below, save your project every so often (Ctrl+Shift+S)*

For Left Ventricle (LV):

- To start segmenting in 3D, you want to view the raw image data while hiding the active selection, which is opaque (i.e. you can’t see through it)
- Hiding the active selection makes it possible to see 1) the outer boundaries of each cardiac chamber and 2) the direction of the cardiac fibers, which will help you define the RV/LV boundary
- Verify that the ‘Show in 3D’ option in the selection window is unchecked
- You can toggle it on/off as needed as you refine the selection
- Adjust the 3D display control as desired
- ‘Volume rendering’ should be checked
- The ‘option’ dropdown menu on the far right gives you 4 different options for 3D display
- *MIP* works best for seeing inside the heart (i.e. reveals the interface between the RV and LV)
- *VRT* works best for seeing surface of heart as you verify how well segmented materials fit together
- Make sure that ‘Orthographic’ view is enabled (not ‘Perspective’ view)
- This view will give you an easier perspective to look at as you rotate the heart before making a selection
- Start by using the 3D viewer by itself (i.e. select the 3D viewer and hide the 2D viewers) and rotate the heart (with the hand icon) to look down the length of the heart (XZ view)
- Click the lasso tool (3^rd^ tab in selection window) and select the ‘Intersect’ option under 3D
- This tool will let you freehand draw in the 3D viewer
- By using the ‘Intersect’ option, the selection you make will be the pixels within your hand drawn selection that are also included in the active selection (i.e. only pixels from “All Signal” material that fall within the freehand drawn selection).
- Be careful when going from the lasso tool selection window back to the 3D viewer; you can very easily click by mistake in the 3D viewer when you intended to rotate with the trackball (hand icon)
- Start by making an outline of the LV larger than it needs to be (i.e. you are selecting beyond the boundaries)
- Look in the 2D viewers to see what areas need to be refined
- For segmenting LV, the XZ and XY views are best
- To pinpoint an area on the 3D viewer, turn on ‘2D crosshairs’ and move the crosshairs to an area of interest on a 2D viewer
- Rotate the heart in the 3D viewer and refine the active selection from this different angle
- Check how it looks against the 2D and 3D image data
- Toggle the ‘Show in 3D’ option on/off to see how the selection looks as you work on it
- Change the 3D display option between MIP and VRT as needed
- Fortunately, you can undo (under Edit menu) if you select within the boundaries of the chamber or click somewhere in the 3D viewer by accident
- Repeat these steps to refine your selection of the outer boundaries of the LV and the RV/LV boundary
- Look at the anatomy of the mouse heart and histology (coronal sections) for further guidance
- Use ‘Fill’ | ‘All slices’ from the Selection dropdown menu, which will include the lumen (for the time being)
- Once you have “carved” the LV as well as you can in 3D, add your selection to a new material labeled “LV”
- For this action (or whenever you reassign pixels to a different material), the lock icon next to the highlighted material must be open to add the actively selected pixels to it
- If there are selected areas seen in a 2D viewer that you cannot easily omit using the 3D lasso method...
- First, add the active selection you already have to the material “LV” and checkmark the ‘2D’ box for “LV”
- On one 2D viewer (don’t switch between them), use the 2D lasso tool and ‘Interpolate’ option in the Selection dropdown menu (shortcut: Ctrl+I) to select areas across multiple slices
- To discard background pixels, highlight “LV” in material list and click the ⊖ button in the selection window to remove the selection entirely, or
- To reassign pixels that will be used for a specific material (e.g. RV), create a new material with that name, highlight it in the material list, and click the ⊕ button in the selection window to reassign the selection to a different material, or
- To retain pixels that will be used for >1 material, highlight “All Signal” in material list and click the ⊕ button in the selection window to send the active selection back
- If there are areas seen in a 2D viewer that you cannot easily include using the 3D lasso method…

*Avoid having to do this by “overselecting” with the 3D lasso (+ ‘Intersect’) as described above*

- First, add the active selection you already have to the material “LV” and checkmark the ‘2D’ boxes for both “LV” and for “All signal”
- On one 2D viewer (don’t switch them them), use the 2D lasso tool and ‘Interpolate’ option in the Selection dropdown menu (shortcut: Ctrl+I) to select areas falling within the “All signal” contours over multiple slices
- Highlight “LV” in material list and click the ⊕ button in the selection window to reassign the active selection to “LV”
- Click on the Magic Wand tool (4^th^ tab in selection window) so that you can reassign pixels in the lumen to a separate material
- Enable ‘All slices’
- Enable ‘Same material only’
- Enable ‘2D preview’
- Checkmark the ‘2D’ box for “LV” so that you can see the outline of the material
- Set the masking range minimum to 0 and adjust the maximum to a level such that the blue pixels only reach the endocardial boundary (i.e. lumen only)
- On a center slice, click within the lumen, which will turn desired pixels purple
- Scroll through the slices and shift-click on any additional unconnected areas within the endocardial boundary that were missed by the magic wand selection
- Create a new material called “LV lumen” to which you can reassign the selected pixels
- Check that the lock icon is open next to “LV” (it is easy to mistake whether it is open or closed)
- Highlight “LV lumen” in material list and click the ⊕ button in the selection window to reassign the active selection to “LV”
- Once finalized, lock the “LV” material (i.e. click the lock icon next to it to close it)
- Now the pixels assigned to “LV” cannot be reassigned unless you unlock the material

For remaining chambers:

- Click the ‘Select’ button to the right of the “All Signal” material
- You should see the 3 remaining chambers, the aortic root, and connective tissue in the 3D viewer
- Repeat the entire process above for RV and the process up until the Magic Wand step for LA and RA
- It will be a faster process for these ones because of the smaller sizes
- For RV, rotating to the short-axis view (XZ) first (as with the LV) and looking from bottom towards the top is probably best
- By keeping “LV” and other fully segmented chambers locked, you can “overselect” in 3D or 2D and be confident that you will not reassign those pixels
- Do not include the aortic root or connective tissue at the top of heart in any selections
- Other options and tools in the Segmentation Editor could also be helpful, such as:
- Grow / Shrink selection options
- Limit line option
- Magic wand tool
- A .labels file (containing the materials) will be created attached to the raw signal data

**Mask generation and chamber wall size quantification**

- Run *Arithmetic* on the .labels file (4 times)
  - Input A: .labels file
  - Input B: same .labels file
  - Results channels: ‘like input A’
  - Write expression as “A==*x*” where *x* is the index of the materials in the .labels file
    - 1 should correspond to “All signal” material (don’t use this one)
    - 2 may correspond to “Inside” (don’t use this one either)
    - 3 should correspond to LV (the 3^rd^ material)
    - 4 should correspond to the 4^rd^ material (i.e., RV)
    - 5 the 5^th^ material (i.e., LA)
    - 6 the 6^th^ material (i.e., RA)
  - Each output generated will be a chamber-specific mask file to be used below for LYVE1 and ER-TR7 signal
- Run *Label analysis* on the .labels file
- A small table will be generated with volume and 3D surface area measurements for each material (i.e. cardiac chamber)
- Use volume or 3D surface area for normalizing other calculated parameters below to the size of individual chambers

**LYVE1 fluorescence signal processing**

Load 16-bit TIFF-OME raw image data for the fluorescence channel that shows lymphatic vessel labeling (e.g. 647 nm)

- In Tcl Console window, type “create HxImageFilters” and hit Enter
- Generates an object “*Wrapper: image filters*”
- Attach this object to the raw LYVE1 signal data
- Use the following parameters:
  - 3D unsharp masking
  - Kernel: 5
  - Sharpness: 0.7
- Run *Interactive TopHat*
  Step 1
- White top-hat
- 3D; connectivity 26
- Kernel size 5
- Click Apply (1/2)

Step 2

- Assess the effect of the TopHat filter by changing Input to Tophat image
- Adjust *Colormap* maximum level down
- Goal is to have low signal intensity of inner muscle, high signal of vessels on periphery of heart, and moderate signal in atrium
- Bright areas corresponding to connective tissue above the ventricles and the aortic root will not be included in the final quantitative results
- Set the thresholding range using *Intensity* *Range,* which will select desired pixels in **blue**
- Maximum level can be 65,000 (upper limit for 16-bit data)
- Minimum level is what needs to be tuned and checked across slices
- Goal is to include the small less intense vessels deep to the surface of the heart but to not include an excessive amount of noise or background signal
- Best to include some excess signal (i.e. set a lower minimum) rather than less (e.g. 1200 instead of 2000)
- Output will be a binary .tophat file (this is the first of two .tophat files)
- Select *Remove Islands* in Segmentation editor
  - Set .filtered image in ‘Image’ dropdown menu
  - Set .tophat image in ‘label field’ dropdown menu
  - From top menu, click Segmentation | *Remove islands….*
  - Opens up a window that allows you to interactively select and remove ‘islands’ – isolated regions of x voxels – keep this window open for steps below
- 15 voxels could be sufficient to remove artifacts / noise
  - Select ‘Apply to: current slice’ (bottom) and then click ‘Highlight all islands’ (takes just a moment to find islands in the current slice)
  - Zoom in (on XY viewer) and review the small highlighted areas carefully by toggling the 2D checkbox for the material on and off
  - Compare the true signal against the selected signal
    - Increase the brightness under ‘Display control’ ‘2D’
  - Increase the voxel size if larger areas need to be removed; decrease if smaller areas
  - Once satisfied, click ‘Apply’ at bottom right and see the effect this has on cleaning up the unimportant areas in the thresholded signal
  - To apply to the remaining slices, use ‘Apply to: all slices’
  - Undo (under Edit menu) may be used to reverse any actions
- If near the top or bottom of the image stack there are extremely bright strips (i.e. a ring of tissue with no underlying signal) you can use the Pick and Move tool (click the 1^st^ tab in the Selection window), deselect ‘all slices’, click inside the strip to select it, and click the ⊖ button to remove it from the .tophat data
- To repeat this process on other slices, you can manually view the other slices and do the same thing OR try select ‘all slices’ and see if this captures what you need to remove (involves some trial and error)
- Once done, save the project to save the modified .tophat file
- Run *Distance Map for Skeleton*
  - Radius 100
- Run *Interactive Top-hat*

Step 1

- Input: .distfield file
- White top-hat
- 3D; connectivity 26
- Kernel size 3
- Click ‘Apply (1/2)’

Step 2

- Adjust intensity range minimum to 1
- Click ‘Apply 2/2’
- Output will be a second .tophat file
  - Only to be used as input for ‘No end map’ option in *Centerline Tree* module (below)
- If not done already, perform segmentation of individual cardiac chambers and run *Arithmetic* on .labels file
  (see **Cardiac chamber segmentation and mask generation** above)
- A .labels file with 4 materials corresponding to the cardiac chambers is made after segmentation
- Running the *Arithmetic* module on each material will generate chamber-specific masks
- These masks are necessary for the subsequent analysis
- Run *Mask* (4 times), once on each mask of the four chambers
- Input: mask file of each chamber
- Input binary image: first .tophat file
- Each output will be a chamber-specific thresholded area for LYVE1 signal
  - Rename these outputs to LV, RV, LA and RA to keep track of the objects
- Run *Centerline Tree* (4 times), once on each output above (LV, RV, LA and RA)
- Input: output from *Mask* of each cardiac chamber
  - Click > symbol to reveal the dropdown menu for ‘No end map’ input
  - Use the second .tophat file as the ‘No end map’ input (this may speed up the processing)
  - Use default tube parameters
  - The ‘number of parts’ is *extremely* important; it restricts the number of possible segments to be found in each chamber
  - You will need to determine empirically what value is reasonable for each chamber
  - LV set to 10,000 for the current study (i.e. P40 hearts)
  - RV set to 7,000
  - LA and RA each set to 3,000
  - Otherwise, you can set this parameter to -1 and the algorithm will continue to run as long as possible
  - In this case, you will need to implement a method for removing extraneous segments (either through image post-processing or filtering out measured values in the next steps)
- Do not create additional objects
- Hold Shift and select all 4 modules, click Apply (this could run for a few hours in the background)
- Run *Spatial graph view* (4 times) on outputs from *Centerline Tree*
- You can visualize average segment radius (referred to as local thickness) or other local parameters
- Run *Spatial graph statistics* (4 times) on outputs from *Centerline Tree*
- This module will generate summary and branch-level measurements, including number of segments and segment radius, length, and tortuosity
- Export the spreadsheets for offline analysis in statistical software

**ER-TR7 fluorescence signal processing**

Load 16-bit TIFF-OME raw image data for the fluorescence channel that shows fibroblasts and reticular fiber labeling

- Run *Structure Enhancement Filter* (SEF) module (based on Frangi filter [1]) on the raw ER-TR7 signal data
- Interpretation type: XY planes
- Tensor type: Hessian
- SD min/max: 1 and 2
- SD step: 1
- Contrast: Bright
- Structure type: Rod
- Run *Convert Image Type*
- 16-bit unsigned
- In Tcl Console window, type “create HxImageFilters” and hit Enter
- Attach this object to the output from previous step
- Run *Grayscale: Sigmoid Intensity Processing* in the module
- Type: 3D
- Alpha: 18,000
- Default beta value
- Minimum: 18,000
- Default maximum value
- Open Segmentation Editor to threshold and refine selection of ER-TR7^+^ fibroblasts and reticular fibers
- Select .filtered image from previous as Image
- Create a new label field and rename it “ER-TR7”
- Click the 8^th^ tab (Top-hat Threshold) in the bottom Selection window

Step 1

- White
- Shape: Cube
- Connectivity: 26
- Kernel size: 2
- Press ‘Compute Top Hat Image’

Step 2

- Open ‘Display control’ window (halfway down Segmentation Editor controls)
- On 2D display, reduce maximum level down to the tail of the histogram (to brighten the display but not oversaturate)
- Set the masking range (highlights masked pixels in **blue**)
- Minimum value: 25% of default
- Default maximum value
- Ensure ‘all slices’ is checkmarked and press ‘Select Masked Voxels’
- Press ⊕ to add selection to ‘Inside’ material
- Rename label file as “fibroblasts”
- Change Image (dropdown menu) to the raw ER-TR7 signal data to do a quality check
- The image may not update automatically; try toggling label field to “NO SOURCE” and then back
- Adjust 2D display control to see raw signal and autofluorescence background
- Check that selected label field is “fibroblasts”
- Scroll through slices to verify that fibroblasts and reticular fibers are fully captured and assigned to the material
- If there are obvious artefacts, you can use the Pick and Move tool (click the 1^st^ tab in the Selection window), deselect ‘All Slices’, click the artefacts, and click the ⊖ to remove from the material
- To repeat this process on other slices, repeat or try selecting ‘All slices’ and see if this cleanly removes what is necessary. This may take some trial and error using ‘Undo’ under Edit menu.
- From top menu, click ‘Segmentation’ and then ‘Remove Islands’
- Opens up a window that allows you to interactively select and remove isolated voxel areas (islands)
- Keep this window open for the steps below
- 15 voxels may be sufficient to remove incorrectly assigned areas
- Select ‘Apply to: current slice’ (bottom) and then click ‘Highlight all islands’ (takes just a moment to find islands in the current slice)
- Zoom in (on XY viewer) and review the small highlighted areas carefully by toggling the 2D checkbox for the material on and off
- Compare the true signal against the selected signal
- Increase the brightness under ‘Display control’
- Increase the voxel size if larger areas need to be removed; decrease if smaller areas
- Once satisfied, click ‘Apply’ at bottom right and see the effect this has on cleaning up the incorrectly assigned areas in the thresholded signal
- To apply to the remaining slices, use ‘Apply to: all slices’
- Undo (under Edit menu) may be used to reverse any actions
- Once done, save project to save the modified material
- If not done already, perform segmentation of individual cardiac chambers and run *Arithmetic* on .labels file
  (see **Cardiac chamber segmentation and mask generation** above)
- A .labels file with 4 materials corresponding to the cardiac chambers is made after segmentation
- Running the *Arithmetic* module on each material will generate chamber-specific masks
- These masks are necessary for the subsequent analysis
- Run *Mask* on each mask of the four chambers
  - Input: mask file of each chamber
  - Input binary image: “fibroblasts” label file object
  - Each output will be a chamber-specific thresholded area for ER-TR7 signal
  - Rename these outputs to LV, RV, LA and RA to keep track of the objects
- Run *Material Statistics* module on each output to determine the ER-TR7 signal volume in each chamber

**Supplementary Figures**

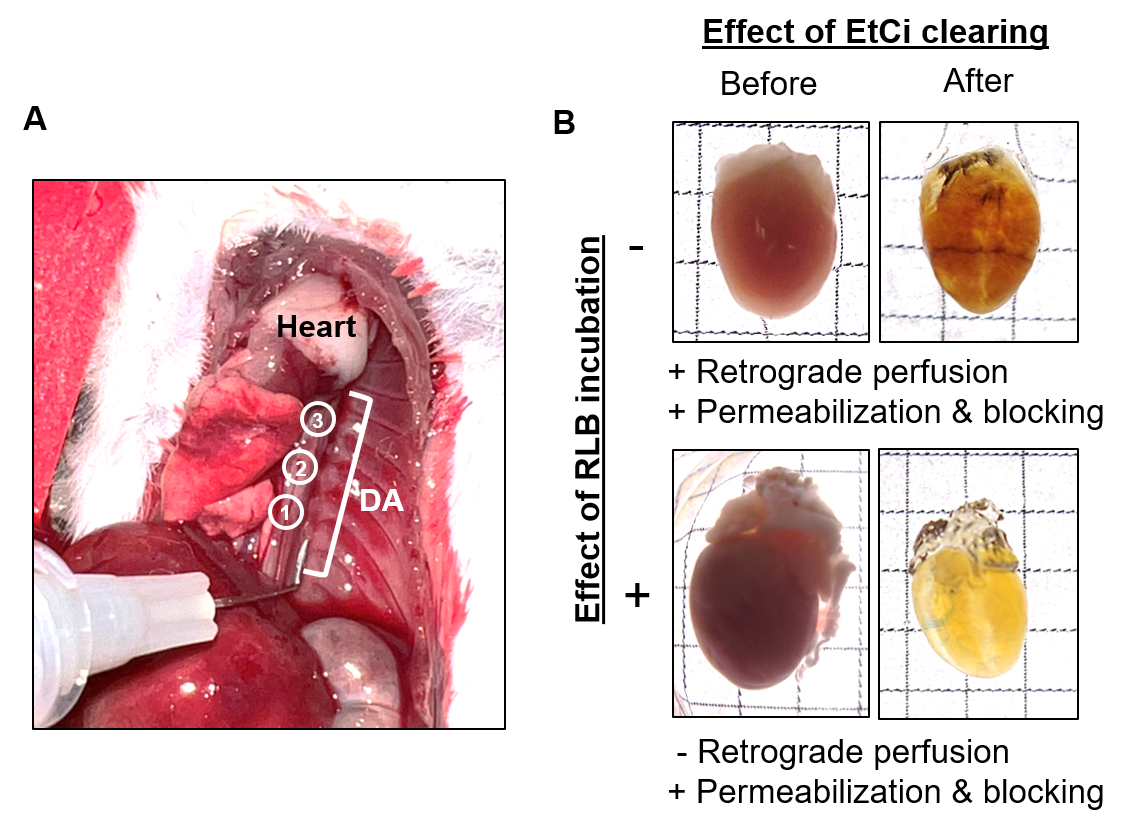

**FIGURE S1** Protocol photos. (A) After inverting lungs to the animal’s right side and cutting the diaphragm, the descending aorta (DA) is visualized prior to *in situ* retrograde perfusion (three injection sites) along the length of the DA with a bent needle and syringe. (B) Application of EtCi-based clearing results in transparency through the depth of the heart (top images) after retrograde perfusion and permeabilization and blocking. Passive incubation in RLB (without prior retrograde perfusion) causes heme elution and subsequent permeabilization, blocking, and EtCi-based clearing leads to transparency and decolorization. Grid dimension is 2.5 mm.

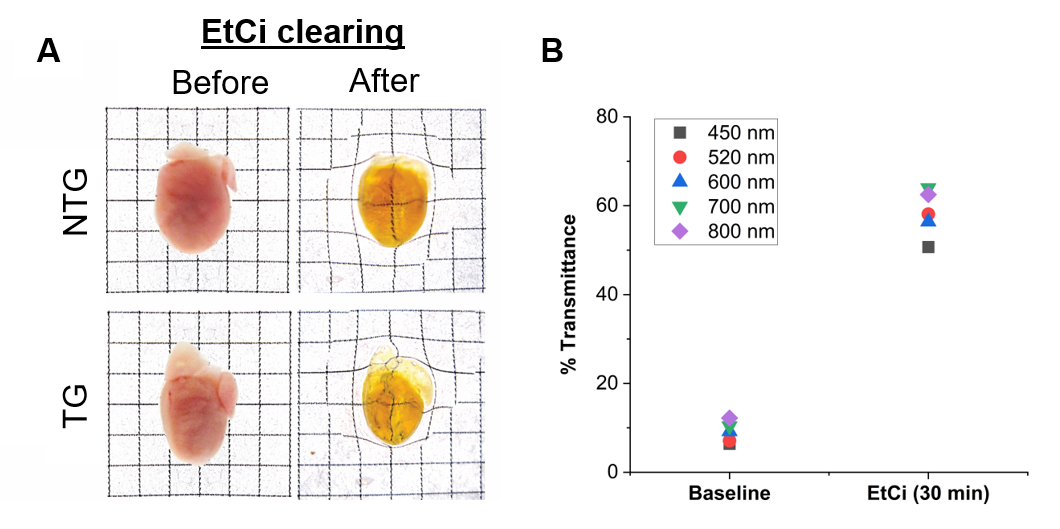

**FIGURE S2** (A) Example photos (anterior perspective) of NTG and TG hearts at start of protocol (after cardiac perfusion and extraction (left)) and by end of protocol (after decolorization, dehydration, and EtCi-based tissue clearing (right)). Hearts on the left are not in a solution; hearts on the right are shown with a drop of EtCi after EtCi-based clearing for 4 hours. Grid dimension is 2.5 mm. (B) To quantify the efficiency of optical tissue clearing in the young adult mouse heart, we compared percent transmittance values (collected with a Jenway model 6300 spectrophotometer) post-fixation and washing to those collected after 30 minutes incubating in EtCi. Laser excitation wavelengths of 450, 520, 600, 700, and 800 nm were used to collect measurements of the percentage of light that passed through a cuvette holding the heart and filled with PBS (baseline) or EtCi (30 min time point).

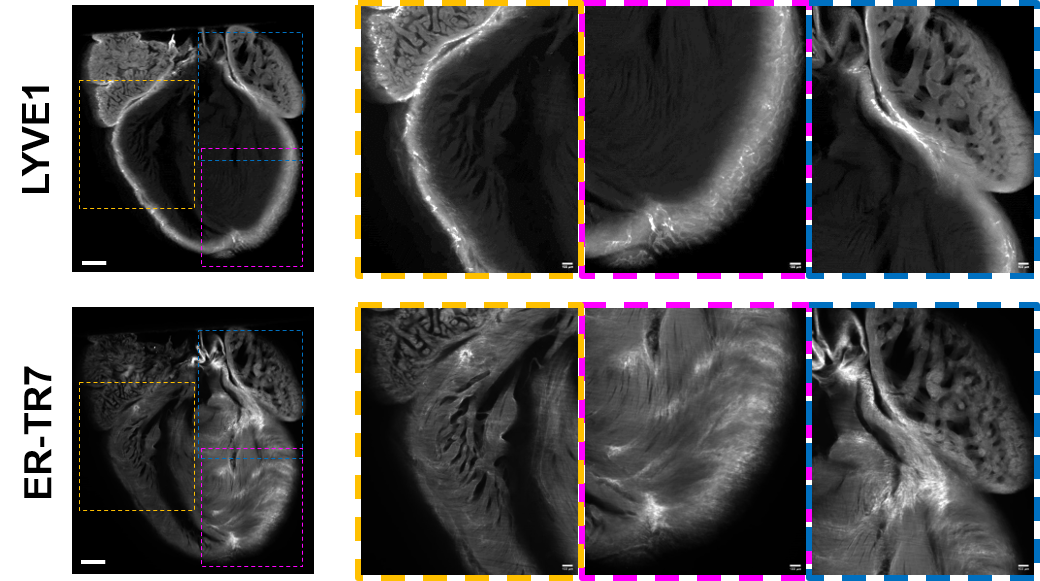

**FIGURE S3** Raw fluorescence signal examples in a TG heart. LYVE1 (top row) and ER-TR7 (bottom row) signals are shown in centerline 2D optical coronal sections (left images). Scale bar: 500 µm. 1.5x-magnified areas from left images are shown on the right: RV (yellow box), LV (magenta box), and left superior (blue box). Scale bar: 100 µm.

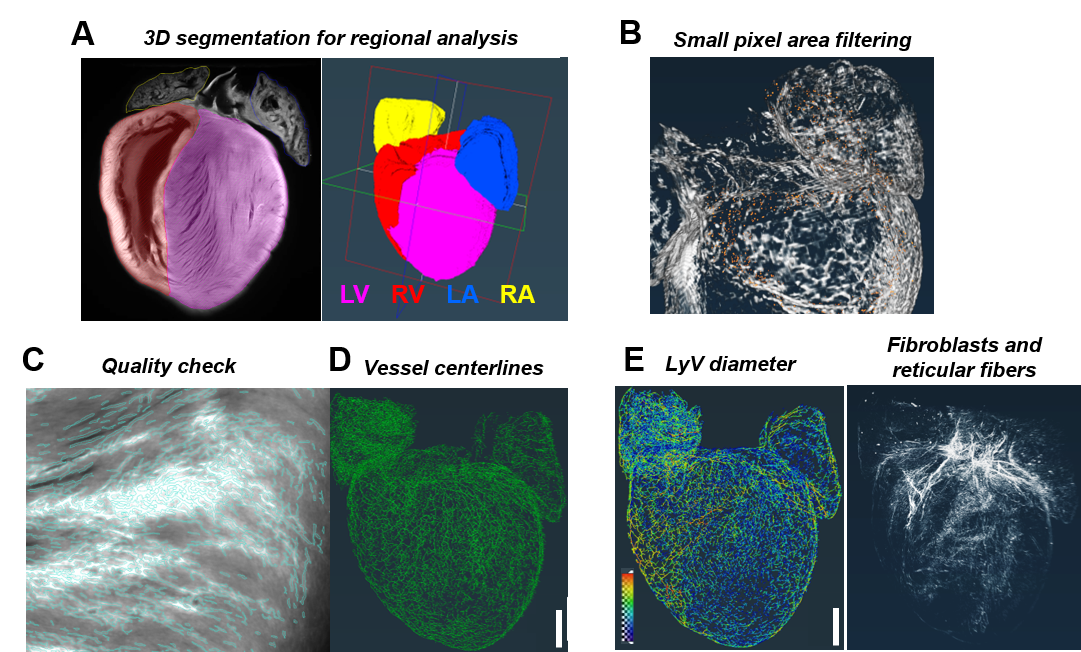

**FIGURE S4** 3D image processing overview using Amira 3D Pro. (A) For efficient manual segmentation of each chamber, the 3D lasso (with Intersect option) was used. For refinement of the segmentations, specific regions seen in virtual 2D view were pinpointed to make adjustments with the 2D lasso. Shown are an optical 2D slice with segmentations drawn over individual chambers (left) and a 3D view of the same heart (right). After excluding the lumens in LV and RV, total volume and 3D surface area of each segmentation were measured. (B) After signal thresholding, small pixel areas (i.e. ‘islands’) that were not filtered out were easily identified and excluded from the true signal by implementing Remove Islands. Shown is a right lateral view of LyVs with small areas (highlighted in red) to be removed. (C) Quality check verification of thresholded signal (cyan outlined areas) matching raw ER-TR7 signal (white intensity areas on underlying grayscale image) in LV. (D) The Centerline Tree module was chosen to identify and trace LyVs given the fact that lymphatic networks have a tree-like structure (no loops) with blunt-ended capillaries. Scale bar: 1 mm. (E) 3D visualization (i.e. spatial graphs) and quantitative data were generated based on (left) regional measurements of LyV parameters and (right) chamber-level measurements of the volume fraction of fibroblasts and reticular fibers. Scale bar: 1 mm.

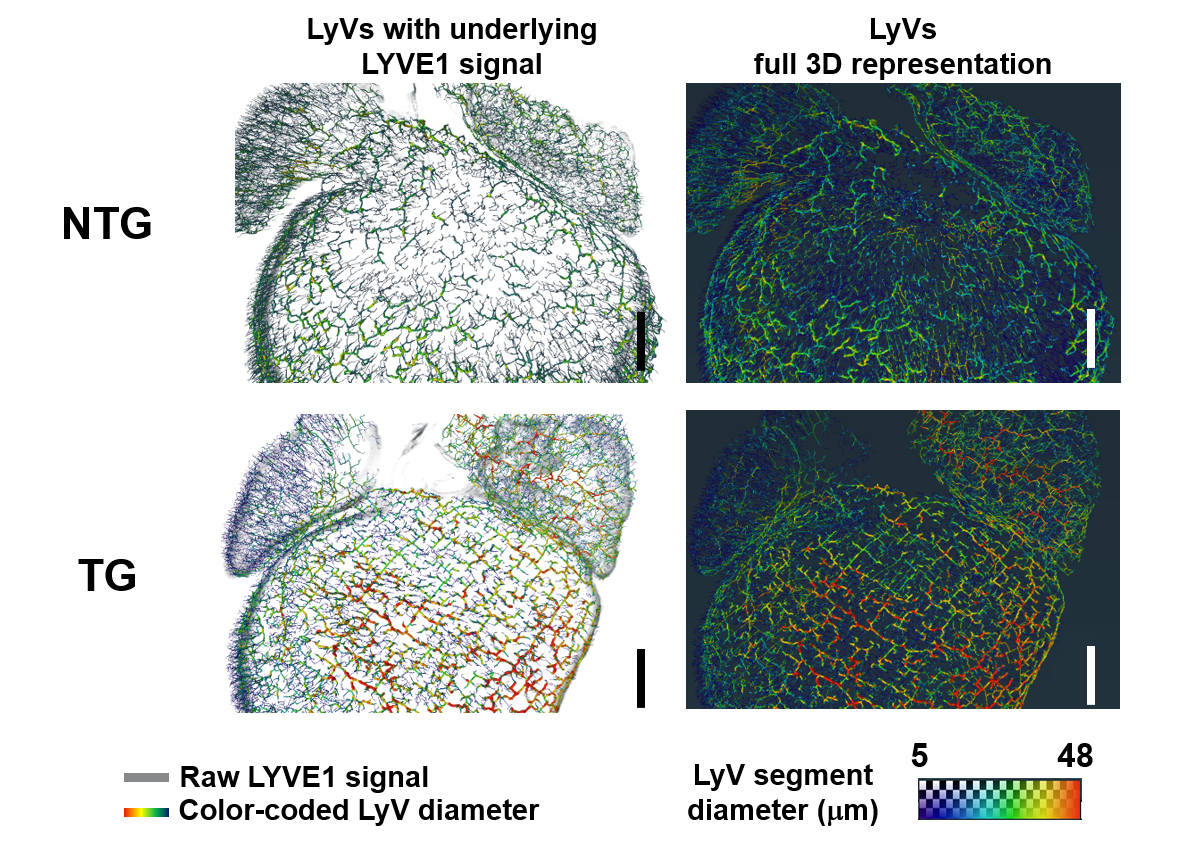

**FIGURE S5** Segmentation and quantitative analysis of LyVs. Left column: 3D images of the posterior halves of an NTG and TG heart with segmented color-coded LyVs (according to their diameters) overlaid on the raw fluorescence signal due to AF647-anti-LYVE1 immunolabeling (grayscale). Right column: Same perspective but showing full 3D representation of LyVs from posterior to anterior (no raw signal). Scale bar: 700 µm.

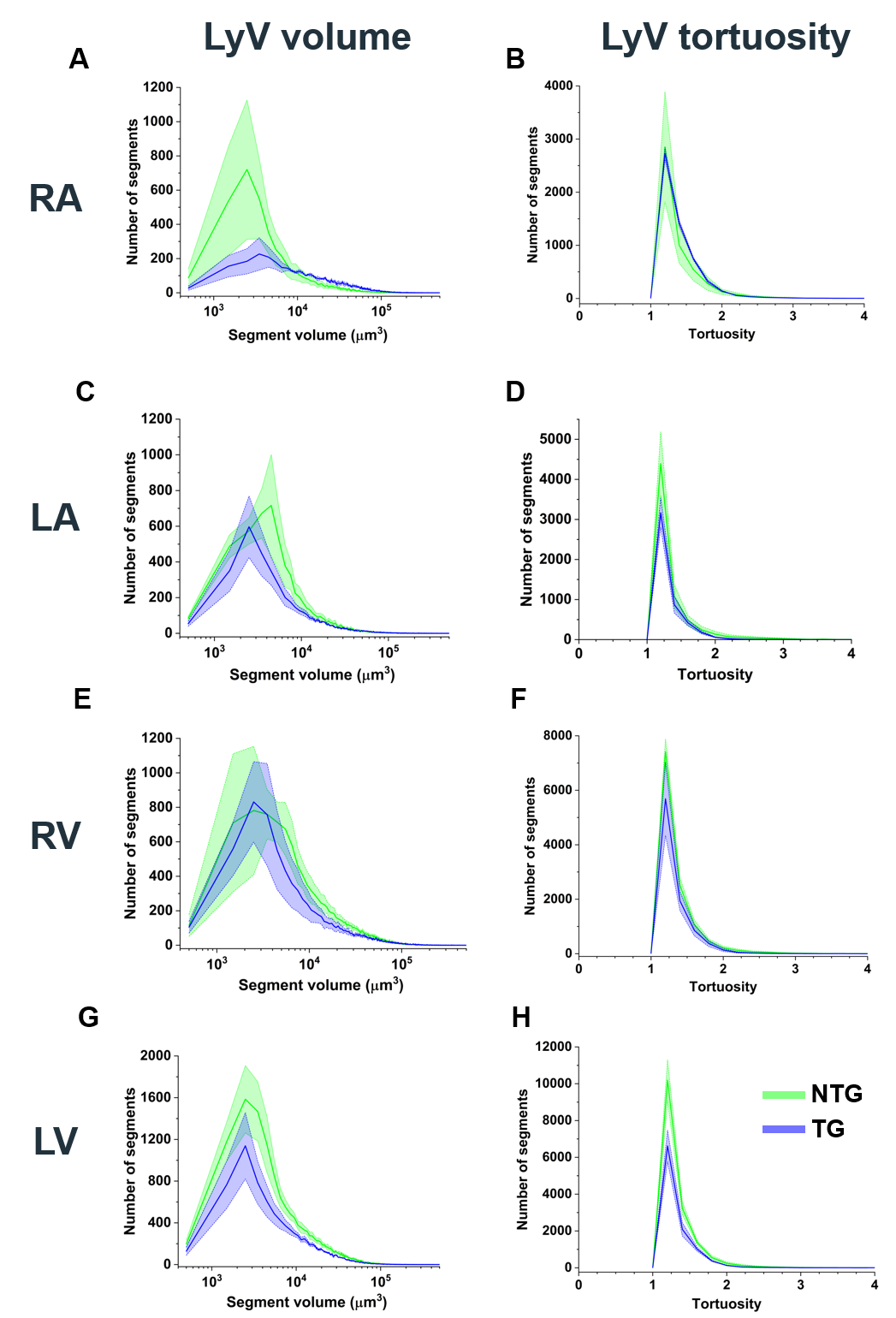

**FIGURE S6** Additional branch-level measurements of LyV segments across chambers. Histograms show distributions of LyV segments for NTG (green) and TG (blue) hearts according to segment volume (left column: A, C, E, G) and segment tortuosity (right column: B, D, F, H) in RA, LA, RV, and LV. Bin sizes: 15000 µm^3^ (volume) and 0.2 (tortuosity). Thick distribution lines: average. Shaded areas: standard error of mean.

**Supplementary Tables**

| Chemicals / Reagents | Stock Concentration or Amount | Final Concentration or Amount | Comments | Manufacturer/  Vendor | Product No. |
| --- | --- | --- | --- | --- | --- |
| **Heparin** | 25 kU | 10 kU / kg body weight | Diluted in PBS | Sigma | H3393-25KU |
| **Phosphate buffered saline without Ca^2+^ / Mg^2+^ (PBS)** | 10X | 1X | For heparin dilution | Sigma | P5493 |
| **Potassium chloride (KCl)** | >/= 99% | 130 mg/mL | Saturated solution in deionized water | Sigma | P3911 |
| **RPMI 1640 medium (RPMI)** |  |  | Contains 10% w/v glycine | Gibco | 11875085 |
| **Red blood cell lysis buffer (RLB)** |  | 1:1 RLB:RPMI |  | Sigma | 11814389001 |
| **Sodium chloride** | >/= 99% | 80 g | For 10X PBS | Sigma | S9888 |
| **Potassium chloride** | >/= 99% | 2 g | For 10X PBS | Sigma | P3911 |
| **Sodium phosphate dibasic** | >99% | 14.4 g | For 10X PBS | Sigma | S9763 |
| **Potassium phosphate monobasic** | >99% | 2.4 g | For 10X PBS | Sigma | P0662 |
| **Sodium hydroxide** | 1.0 N |  | For pH adjustment | Sigma | S2770 |
| **Paraformaldehyde (PFA)** | 16% | 2% | Diluted in PBS | EMS | 15710-S |
| **Triton X-100** |  | 0.3% v/v | Diluted in RPMI | Sigma | T8787 |
| **Bovine serum albumin (BSA)** |  | 1% w/v | Diluted in RPMI | Sigma | A2058 |
| **Dimethylformamide (DMF)** | 99% | stock concentration | For dye preparation | Sigma | D4551 |
| **DyLight 550 NHS ester (DyL550)** | 1 mg | 10 mg/mL in DMF | Stock preparation | Fisher | 62262 |
| **Ethanol (EtOH)** | 99% | 50%, 70%, 99% | Diluted in deionized water | Sigma | 24102 |
| **Tetrahydrofuran (THF)** | 99% | stock concentration | For TEA dilution | Sigma | 401757 |
| **Triethylamine (TEA)** | 99% | 1:10 in THF | Drop-wise addition for pH adjustment | Sigma | T0886 |
| **Ethyl cinnamate (EtCi)** | 99% | stock concentration | Solvent with high refractive index (RI) to match  RI of tissue | Sigma | 112372 |
| **α-thioglycerol (TG)** | ≥97% | 30 µL/10 mL EtCi | To help prevent gradual loss of tissue transparency | Sigma | M1753 |
| Supplies | Specifications | | Comments | Manufacturer/ Vendor | Product No. |
| **Isoflurane** |  | | Inhaled anesthetic | Henry Schein | 1182098 |
| **Lab tape** |  | | To secure mouse paws |  |  |
| **Alcohol wipes** |  | | To flatten hair on mouse ventrum | BD | 326895 |
| **Syringes** | 5 mL Luer-Lok | | For perfusion | Fisher | 14-829-45 |
| **Gauze pads** |  | |  |  |  |
| **Insulin syringes** | 0.5 mL, 29G1/2 | | For heparin | BD | 305930 |
| **Beveled needles** | 27G1/2 | | For perfusion; For pinning paws after thoracotomy | Fisher | 14-826-48 |
| **Cotton tip applicators** |  |  |  |  |  |
| **Wax-lined dissecting tray** |  |  |  | Cole Parmer | EW-10915-14 |
| **Sharp-pointed dissecting scissors** |  |  |  | Fisher | 19-027493 |
| **Straight Vanna Scissors** |  |  | Appropriate for microdissection | Fine Surgical Tools | 91500-09 |
| **Curved Graefe forceps** |  |  | Appropriate for handling mouse tissue | Fine Surgical Tools | 11051-10 |
| **Needle holders** |  |  | Appropriate for clamping a needle during perfusion | Fine Surgical Tools | 12003-15 |
| **Straight ultrafine point tweezers** |  |  | Appropriate for picking up heart by aortic root | Fisher | 12-000-122 |
| **Untreated plastic**  **well plates** | 6-well |  | For washing and fixation after extraction | Fisher | 07-201-588 |
|  | 24-well |  | For incubation steps (except dehydration and tissue clearing) | Fisher | 07-201-590 |
| **Aluminum foil** |  |  | For light protection of samples |  |  |
| **Parafilm** |  |  | To seal well plate | Fisher | 13-374-12 |
| **Slide-A-Lyzer dialysis cassettes** | 0.1-0.5 mL, 10,000 MW cutoff | | For dialyzing antibody (max volume 0.5 mL) after conjugation to fluorescent dye | Fisher | 66383 |
| **Slide-A-Lyzer syringes with 18G needles** | 1 mL Luer-Lok |  | For loading cassettes | Fisher | PI66494 |
| **Nalgene polypropylene beaker** | 2 L |  | For dialysis in PBS | Fisher | 02-591-10H |
| **Magnetic stir bars** |  |  |  | Fisher | 14-512-124 |
| **Microtubes** | 5 mL |  | For dehydration | Fisher | 03-391-270 |
| **Glass vials with caps** | 20 mL |  | For tissue clearing | Fisher | 0334025P |
| IgG antibody | Stock Conc. | Working Dilution | Comments | Manufacturer | Catalog No. |
| **goat anti-mouse LYVE1** | 0.2 mg/mL | 1:50 | Unconjugated primary antibody | R&D Systems | AF2125 |
| **Alexa Fluor 647 (AF647) AffiniPure donkey anti-goat (H+L)** | 1.5 mg/mL | 1:250 | Fluorescent secondary antibody | Jackson Immuno-research | 705-605-147 |
| **rat anti-mouse ER-TR7** | 1.5 mg /mL | 1:25 | Unconjugated primary antibody | BioXCell;  (Bio-Rad) | Discontinued  (Bio-Rad MCA2402 is now available) |
| Equipment | Part or Catalog No. | | Comments | Manufacturer/ Supplier |  |
| **Isoflurane vaporizer and inhalation tubing** | VetFlo-1205S | | 3-5% flow rate to induce deep anesthesia | Kent Scientific |  |
| **Gooseneck stereomicroscope with ring light** | SE508-FRL | | For *in situ* perfusion and microdissection | Amscope |  |
| **Precision scale** |  | | For mouse body and heart weights |  |  |
| **Magnetic stir plate** | 97048-754 | | For antibody dialysis | VWR |  |
| **Water purification system** |  | | To obtain deionized water |  |  |
| **3D shakers with dimpled mat** | 10034-220 | | For washing, incubation, dehydration, and tissue clearing steps | VWR |  |
| **pH meter** |  |  | For pH adjustment |  |  |
| **UltraMicroscope II with Zoom body or Super Plan configuration** |  |  | LSFM (emission detection above 561 nm in two or three channels) | Miltenyi Biotec |  |
| **Amira 3D Pro** | v2021.1 |  | 3D image visualization and analysis software | FEI Company |  |

**TABLE S1** Checklist of materials used in this study and suggested for protocol users

| GENOTYPE | HW  (g) | BW  (g) | TL  (mm) | HW/BW (x 10^-3^) | HW/TL  (x 10^-3^) |
| --- | --- | --- | --- | --- | --- |
| NTG | **0.0806** | **20.26** | **1.577** | **3.98** | **51.1** |
| NTG | **0.0834** | **20.03** | **1.441** | **4.16** | **57.9** |
| NTG | **0.1267** | **24.31** | **2.038** | **5.21** | **62.2** |
| TG | **0.0696** | **19.24** | **1.592** | **3.62** | **43.7** |
| TG | **0.0853** | **19.83** | **1.855** | **4.30** | **46.0** |
| TG | **0.0895** | **24.04** | **1.713** | **3.72** | **52.2** |

**TABLE S2** General characteristics of P40 mice. Heart weight (HW), body weight (BW), and right tibia length (TL) were recorded for the P40 NTG and TG mice analyzed (n=3/group). HW/BW and HW/TL ratios were then calculated.

**Supplementary Video (see video file)**

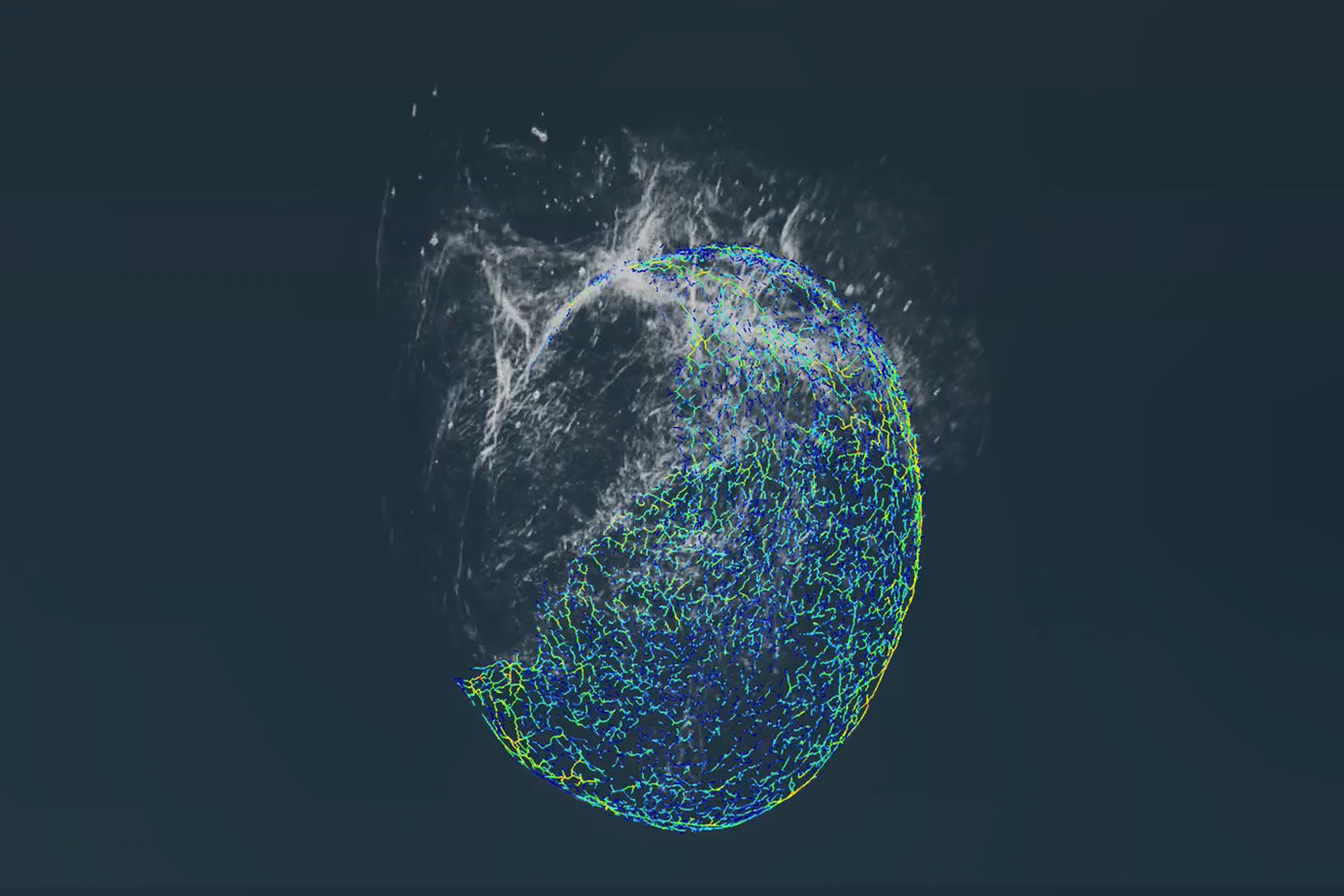

**VIDEO S1** Rotating illustration of the 3D rendering of fibroblasts and reticular fibers throughout a TG heart as well as the distribution and size of LyVs in each cardiac chamber.
